# Neutrophil adhesion in brain capillaries contributes to cortical blood flow decreases and impaired memory function in a mouse model of Alzheimer’s disease

**DOI:** 10.1101/226886

**Authors:** Jean C. Cruz Hernández, Oliver Bracko, Calvin J. Kersbergen, Victorine Muse, Mohammad Haft-Javaherian, Maxime Berg, Laibaik Park, Lindsay K. Vinarcsik, Iryna Ivasyk, Yiming Kang, Marta Cortes-Canteli, Myriam Peyrounette, Vincent Doyeux, Amy Smith, Joan Zhou, Gabriel Otte, Jeffrey D. Beverly, Elizabeth Davenport, Yohan Davit, Sidney Strickland, Costantino Iadecola, Sylvie Lorthois, Nozomi Nishimura, Chris B. Schaffer

## Abstract

The existence of cerebral blood flow (CBF) reductions in Alzheimer’s disease (AD) patients and related mouse models has been known for decades, but the underlying mechanisms and the resulting impacts on cognitive function and AD pathogenesis remain poorly understood. In the APP/PS1 mouse model of AD we found that an increased number of cortical capillaries had stalled blood flow as compared to wildtype animals, largely due to leukocytes that adhered in capillary segments and blocked blood flow. These capillary stalls were an early feature of disease development, appearing before amyloid deposits. Administration of antibodies against the neutrophil marker Ly6G reduced the number of stalled capillaries, leading to an immediate increase in CBF and to rapidly improved performance in spatial and working memory tasks. Our work has thus identified a cellular mechanism that explains the majority of the CBF reduction seen in a mouse model of AD and has also demonstrated that improving CBF rapidly improved short-term memory function. Restoring cerebral perfusion by preventing the leukocyte adhesion that plugs capillaries may provide a novel strategy for improving cognition in AD patients.

## Introduction

Alzheimer’s disease (AD) is the most common form of dementia in the elderly, worldwide. AD is characterized by a rapid and progressive cognitive decline accompanied by several pathological features, such as the accumulation of amyloid-beta (Aβ) plaques in brain tissue and along blood vessels as cerebral amyloid angiopathy, the hyperphosphorylation of tau proteins and formation of neurofibrillary tangles in neurons, increased density and activation of inflammatory cells, and ultimately the death of neurons and other brain cells^1^.

Vascular dysfunction has been strongly implicated in the pathogenesis of AD. Many of the primary risk factors for AD are associated with compromised vascular structure and function, such as obesity, diabetes, atherosclerosis, and hypertension^2^. Brain blood flow is also severely compromised in AD, with both patients with AD^3-5^ and mouse models of AD^6-8^, which overexpress mutated forms of amyloid precursor protein (APP), exhibiting cortical cerebral blood flow (cCBF) reductions of ~25% early in disease development. Several mechanisms for this hypoperfusion had been proposed including constriction of brain arterioles^9^, loss of vascular density*^10^*, and changes in neural activity patterns and/or in neurovascular coupling*^11,12^*, but a full understanding of the underlying mechanisms for CBF reduction in AD has not emerged.

These large blood flow decreases could contribute to the cognitive symptoms of AD and drive disease progression. Cognitive functions, such as attention, were immediately impaired by CBF reductions of ~20% in healthy humans*^13^*. When CBF was chronically reduced by ~35% in wildtype (wt) mice, spatial memory deficits were observed, accompanied by pathological changes in the brain including increased inflammation*^14^*. In addition, impairing blood flow in AD mouse models led to an increase in Aβ deposition, suggesting that blood flow deficits can worsen Aβ pathology^14,15^. These data suggest that the decreased CBF in AD likely contributes to both the cognitive dysfunction and to disease progression.

Because CBF reductions have been a recognized and important aspect of AD, yet have not been well explained, we sought to uncover the cellular basis for these flow reductions in APP/PS1 mice.

## Results

To investigate cortical hypoperfusion in AD, we used *in vivo* two-photon excited fluorescence (2PEF) microscopy to image the cortical vasculature in APP/PS1 mice*^16^* (Fig. 1a) and looked for occluded vessels (Fig. 1b). We observed no obstructions in arterioles or venules, but APP/PS1 mice had a 4-fold elevation in the number of capillaries with stalled blood flow as compared to age- and sex-matched, wt littermates (Fig. 1c, movie S1 and S2). The number of stalled capillaries was elevated by 12 weeks of age in APP/PS1 mice, and remained elevated throughout disease progression (Fig. 1d). Flowing and stalled capillaries (Fig. 1e) had about the same distance distribution relative to the nearest penetrating arteriole (Fig. 1f) or ascending venule (Fig. 1g). The incidence of capillary stalling did not increase with Aβ plaque density (Extended Data Fig. 1), and was the same in awake and anesthetized animals (movie S3 and S4; Extended Data Fig. 2). Capillary stalling was similarly elevated in TgCRND8 mice, a different mouse model of APP overexpression (Extended Data Fig. 3)*^17^*, suggesting that this capillary plugging phenomena is not unique to the APP/PS1 mouse model.

**Fig. 1.**
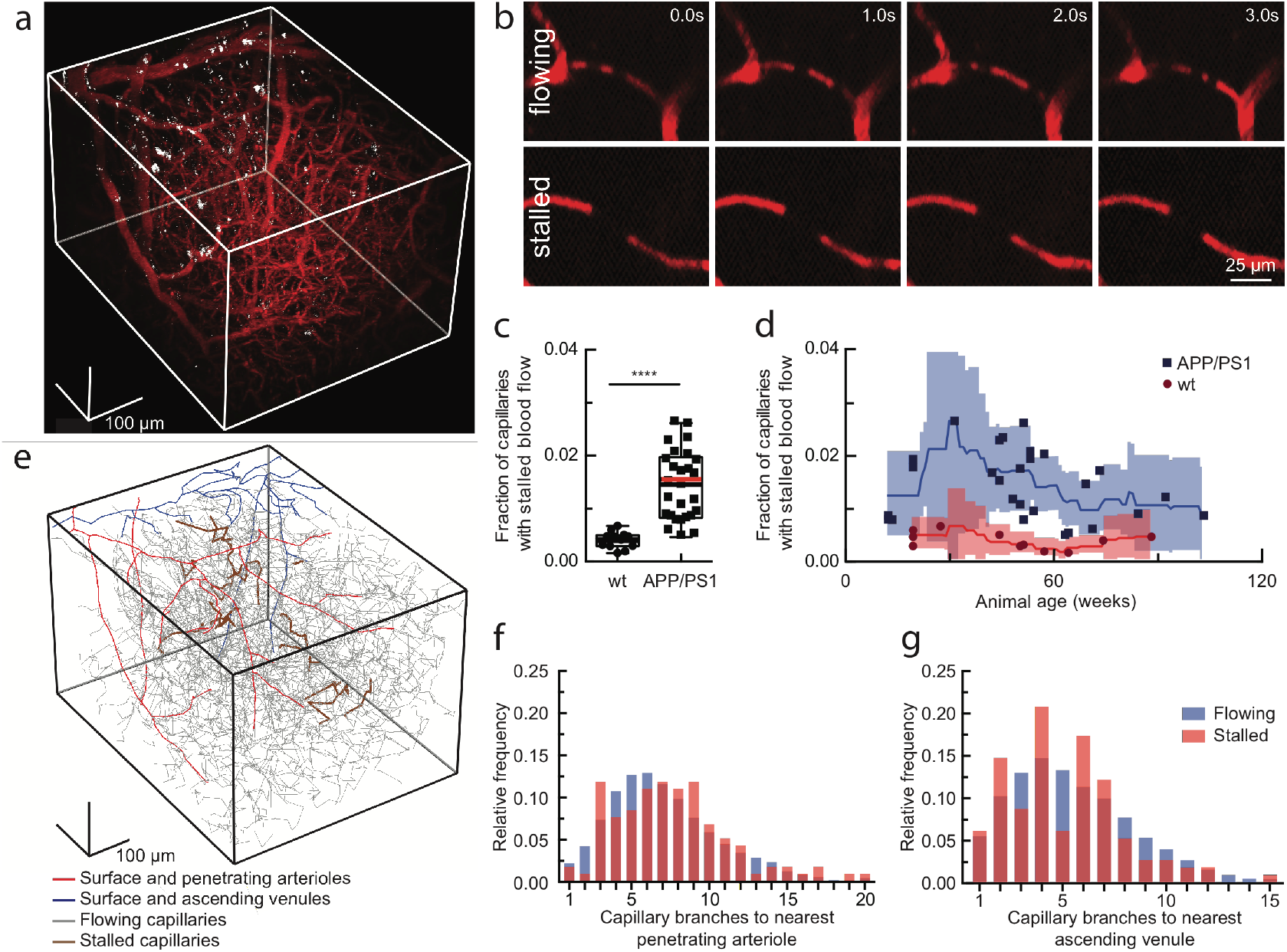
2PEF imaging of mouse cortical vasculature revealed a higher fraction of plugged capillaries in APP/PS1 mice. (a) Rendering of 2PEF image stack of cortical vasculature (red; Texas Red dextran) and amyloid deposits (white; methoxy-X04). (b) Individual brain capillaries were scored as flowing or stalled based on the motion of unlabeled blood cells (black) within the fluorescently labeled blood plasma (red), (c) Fraction of capillaries with stalled blood flow in APP/PS1 and wt mice; (d) same data as a function of age. One outlier not shown: APP/PS1, 42 weeks, 4.4% stalled, (e) Tracing of the vascular network in panel A, with stalled capillaries indicated in brown, (f) and (g) Histograms showing the topological location of flowing and stalled capillaries relative to the nearest penetrating arteriole and ascending venule, respectively.

Using labeling strategies to distinguish leukocytes, platelets, and RBCs (Fig. 2a), we found the majority of stalled capillary segments contained a leukocyte (Fig. 2b). Stalled capillaries had a modestly smaller average diameter than flowing capillaries (Fig. 2c), but no difference in the density of nearby Aβ deposits (Fig. 2d). Most plugged capillaries were transiently stalled with a half-life of less than 5 min, while one-third remained stalled for 15 min and 10% began flowing and then re-stalled within 15 min (Fig. 2e; Extended Data Fig. 4). We also observed that some capillary segments alternated between flowing and stalled in repeated imaging sessions over weeks (Fig. 2f). The same capillaries were stalled across multiple imaging sessions about ten times as frequently as predicted by a statistical model that assumed each capillary had an equal probability of being stalled at any time point (Fig. 2g). Taken together, these data suggest that the capillary stalls were caused by leukocytes adhering in a distinct subset of capillary segments.

**Fig. 2.**
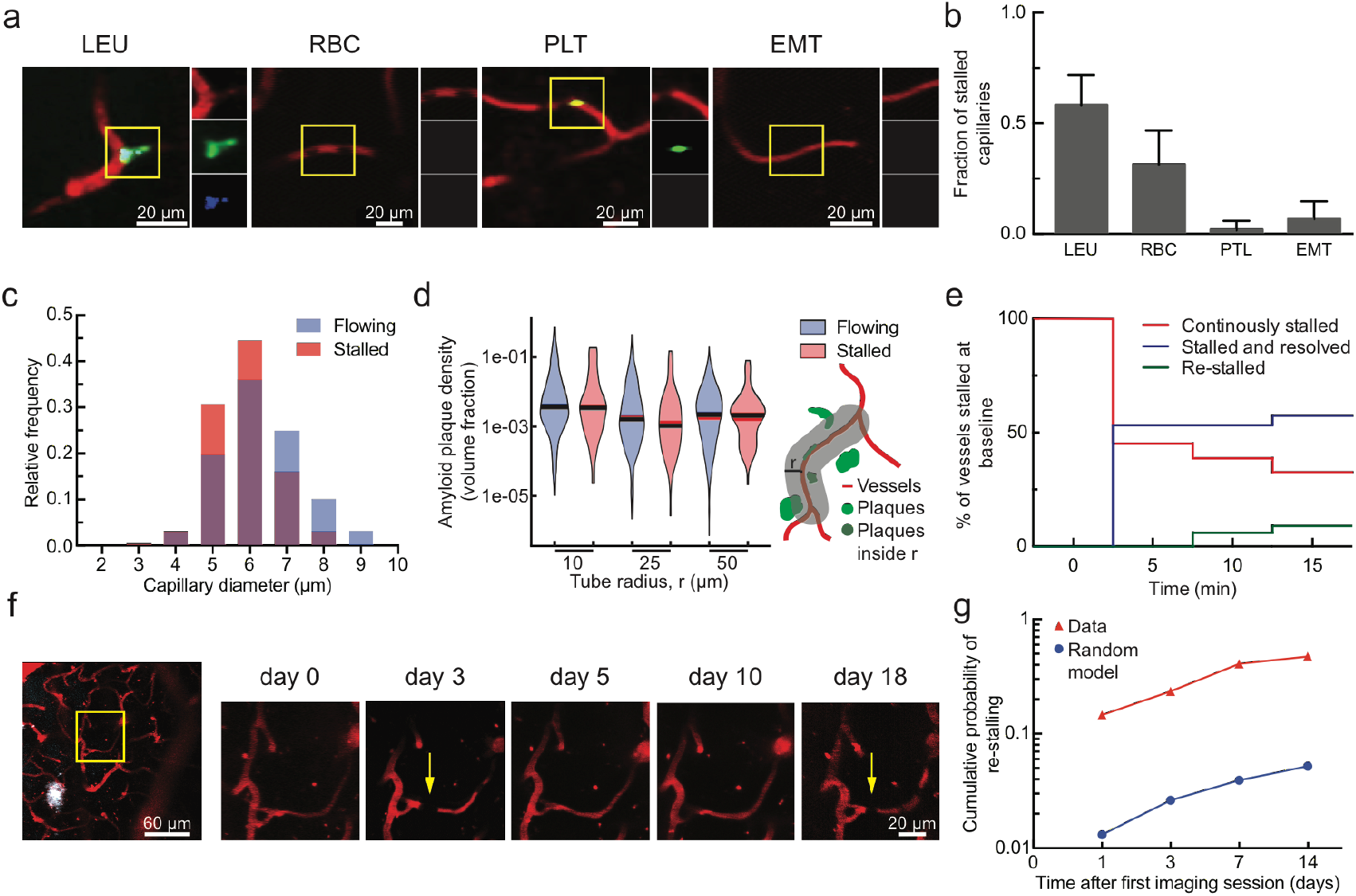
Characterization of the cause, location, and dynamics of capillary occlusions in APP/PS1 mice. (a) 2PEF images of capillaries stalled by a leukocyte (LEU), RBCs, and platelet aggregates (PLT), or found empty of blood cells (EMT), distinguished by fluorescent labels (red: Texas Red-labeled blood plasma; green: rhodamine 6G-labeled leukocytes and platelets; blue: Hoechst-labeled leukocyte nuclei), (b) Fraction of stalled capillaries stalled by LEU, RBC, PLT, or EMT. (c) Histogram of the diameter of flowing and stalled capillaries (Averages: 5.8±0.05 μm (stalled), 6.3±1.1 μm (flowing); p<0.0001). (d) Violin plot of the density of amyloid deposits within tubes of different radii that followed the capillary centerline for flowing and stalled capillary segments, (e) Fraction of stalled capillaries that remained stalled (red), resumed flowing (green), or resumed flowing and then re-stalled (blue) over 15 minutes, (f) 2PEF images of the same capillary alternately stalled (arrows) and flowing (white: methoxy-X04). (g) Cumulative probability of an initially stalled capillary to be observed stalled again at any subsequent imaging time point, showing both real observations and predictions from a model that assumed each capillary had an equal probability of stalling at each time point.

To determine the class of leukocyte causing the capillary stalls, we injected fluorescently-labeled antibodies against Ly6G (α-Ly6G; 4 mg/kg animal weight), a neutrophil surface marker, and serendipitously found that this reduced the number of stalled capillaries within 10 min (Fig. 3a and Extended Data Fig. 5). Isotype control (Iso-Ctr) antibodies did not impact capillary stalling. The rapid resolution of the capillary stalls suggests that α-Ly6G interfered with leukocyte adhesion. Median volumetric blood flow in penetrating arterioles, measured using 2PEF (Fig. 3b), increased by 26% in young (3-4 months) and 32% in aged (11-14 months) APP/PS1 mice one hour after α-Ly6G administration (Fig. 3c). This increase in blood flow was due to an increase in RBC speed and not an increase in vessel diameter (Extended Data Fig. 6). Penetrating arterioles with lower baseline flow tended to show larger flow increases (Extended Data Fig. 7). Iso-Ctr antibodies did not change blood flow in APP/PS1 mice, nor did α-Ly6G in wt animals (Fig. 3c). Further, capillary stalling decreased and penetrating arteriole flow increased one day after administration of antibodies against LFA-1, which depleted circulating leukocytes (Extended Data Fig. 8). Across all antibody and control treatments, penetrating arteriole flows increased (decreased) when the number of stalled capillaries decreased (increased) (Extended Data Fig. 9). We also used arterial spin labeled MRI (ASL-MRI) to measure cortical cerebral blood flow (cCBF) in 7-9-month old animals (Fig 3d). At baseline, average cCBF in APP/PS1 mice was 17% lower than in wt animals (Fig. 3e). cCBF increased by 13% in APP/PS1 mice at ~5 hr after α-Ly6G administration, recovering about two-thirds of the deficit relative to wt animals, but was unchanged in APP/PS1 mice given Iso-Ctr antibodies or wt mice given α-Ly6G (Fig. 3e). Thus, administration of α-Ly6G led to a rapid reduction in the number of capillary stalls that was accompanied by an increase in cCBF in APP/PS1 mice. The specificity of Ly6G expression suggests that the cell type responsible for the capillary stalls was neutrophils^18^.

**Fig. 3.**
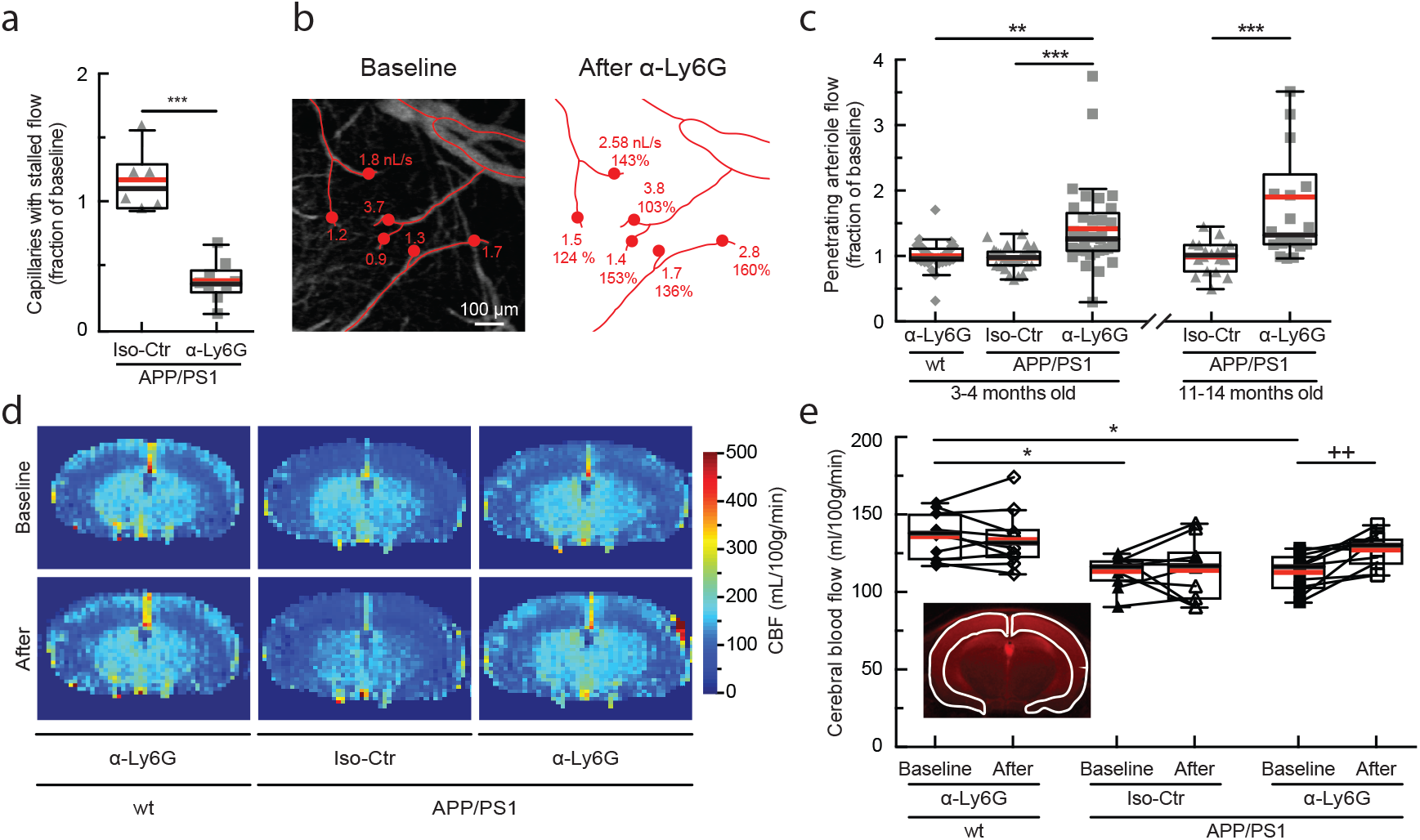
Administration of antibodies against Ly6G reduced the number of stalled capillaries and increased cCBF in APP/PS1 mice. (a) Number of capillaries with stalled blood flow ~1 hr after α-Ly6G or Iso-Ctr antibody administration shown as a fraction of the number of stalled capillaries at baseline. (b) Projection of 2PEF image stack of brain surface vasculature, with surface (red lines) and penetrating (red dots) arterioles identified. For each penetrating arteriole, volumetric blood flow is indicated at baseline (left) and after α-Ly6G administration (right), along with the percentage of baseline flow. (c) Volumetric blood flow in penetrating arterioles measured 60-90 min after α-Ly6G or Iso-Ctr antibody administration in young and old APP/PS1 mice and wt control animals shown as a fraction of baseline arteriole flow. (d) CBF map measured using ASL-MRI at baseline and ~5 hr after administration of α-Ly6G or Iso-Ctr antibodies in APP/PS1 and wt mice. (e) cCBF measurements (ASL-MRI, inset indicates ROI on T2 MRI image) at baseline and ~5 hr after administration of α-Ly6G or Iso-Ctr antibodies in APP/PS1 and wt mice.

We next tested whether α-Ly6G administration improves cognitive function in APP/PS1 mice (Fig. 4a). In the object replacement (OR) test of spatial short-term memory (Fig. 4b), a single dose of α-Ly6G in ~11-month old APP/PS1 mice improved performance to the level of wt animals at 3 and 24 hours after administration (Fig. 4c; Extended Data Fig. 10). APP/PS1 mice treated with Iso-Ctr antibodies showed no change, nor did wt animals with α-Ly6G (Fig. 4c). Similarly, α-Ly6G improved performance of APP/PS1 mice in the Y-maze test of working memory (Fig. 4d; Extended Data Fig. 11). We detected no improvement in sensory-motor function (balance beam walk, (Extended Data Fig. 12) and a trend toward reduced depression- and anxiety-like behavior (forced swim, Extended Data Fig. 13) in APP/PS1 mice with α-Ly6G. We continued to treat these animals with α-Ly6G every three days for a month, resulting in depletion of neutrophils (Extended Data Fig. 14). After this regimen, APP/PS1 mice exhibited short-term memory performance that matched wt animals in OR (Fig. 4c), Y-maze (Fig. 4d), and novel object recognition (NOR) (Fig. 4e; Extended Data Fig. 16). We saw no improvement in sensory-motor function (Extended Data Fig. 12) and a trend toward decreased depression- and anxiety-like behavior (Extended Data Fig. 13).

**Fig. 4.**
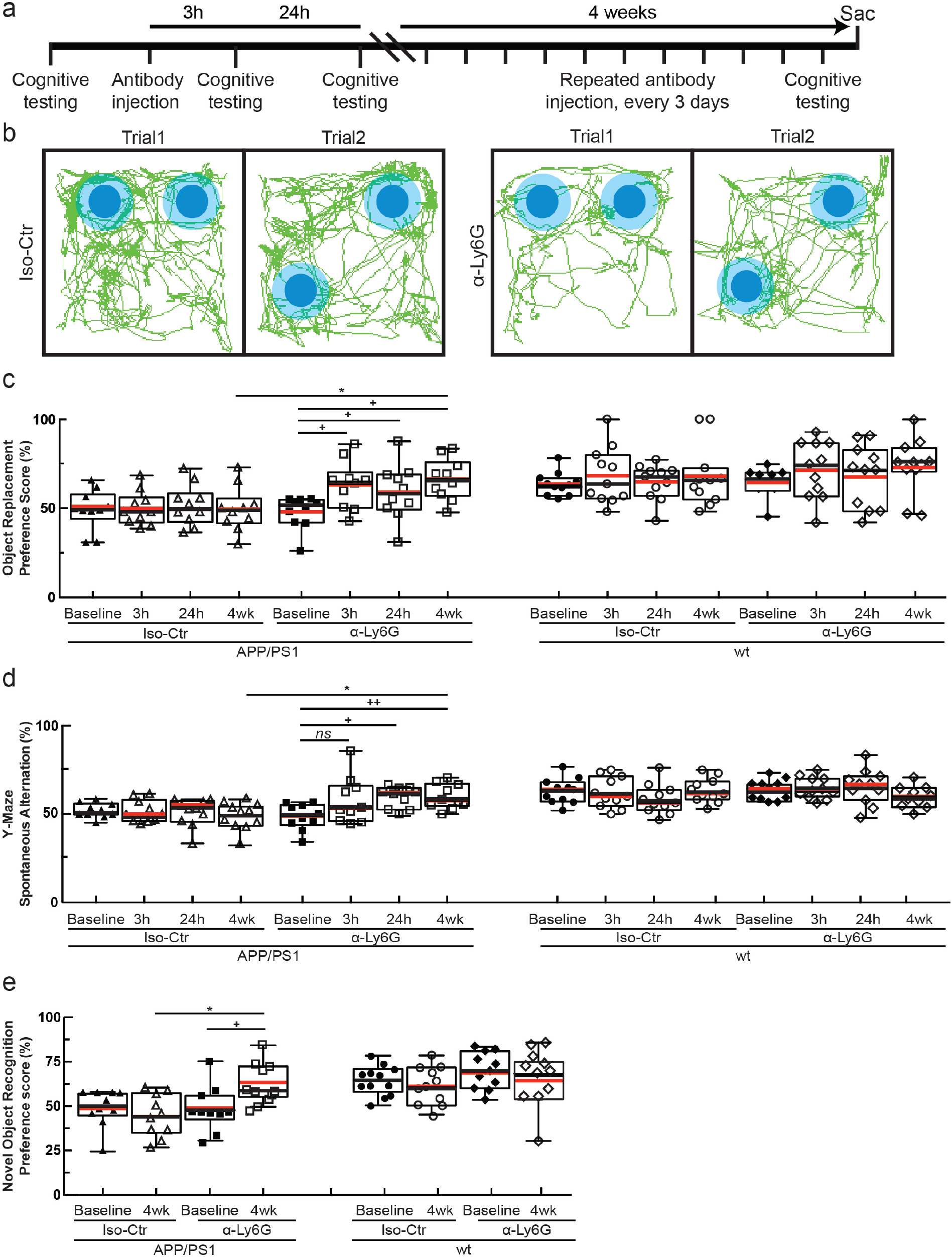
Administration of α-Ly6G improved short-term memory. (a) Experimental timeline for behavioral studies. (b) Tracking of mouse nose location from video recording during training and trial phases of OR task taken 3-5 hr after administration of α-Ly6G or Iso-Ctr antibodies in APP/PS1 mice. (c) Preference score in OR task and (d) spontaneous alternation in Y-maze task for APP/PS1 and wt mice at baseline and at 3 hr and 24 hr after a single administration of α-Ly6G or Iso-Ctr antibodies, and after 4 weeks of treatment every three days. (e) Preference score in NOR task for APP/PS1 and wt mice at baseline and after 4 weeks of treatment every three days.

Because one of the clearance pathways for Aβ is through the vasculature*^19^* we assessed whether improving cCBF with α-Ly6G decreases the concentration of Aβ monomers and aggregates. Using enzyme-linked immunosorbent assays (ELISAs) of brain extracts from the animals that received one month of antibody treatment, we found that α-Ly6G reduced the concentration of Aβ_1-40_ compared to Iso-Ctr antibodies (Fig. 5a), while the concentration of Aβ_1-42_ (Fig. 5b) and aggregates of Aβ (Extended Data Fig. 16d) remained unchanged. We saw no difference in the number and density of Aβ plaques between α-Ly6G and Iso-Ctr treated animals (Extended Data Fig. 16).

**Fig. 5.**
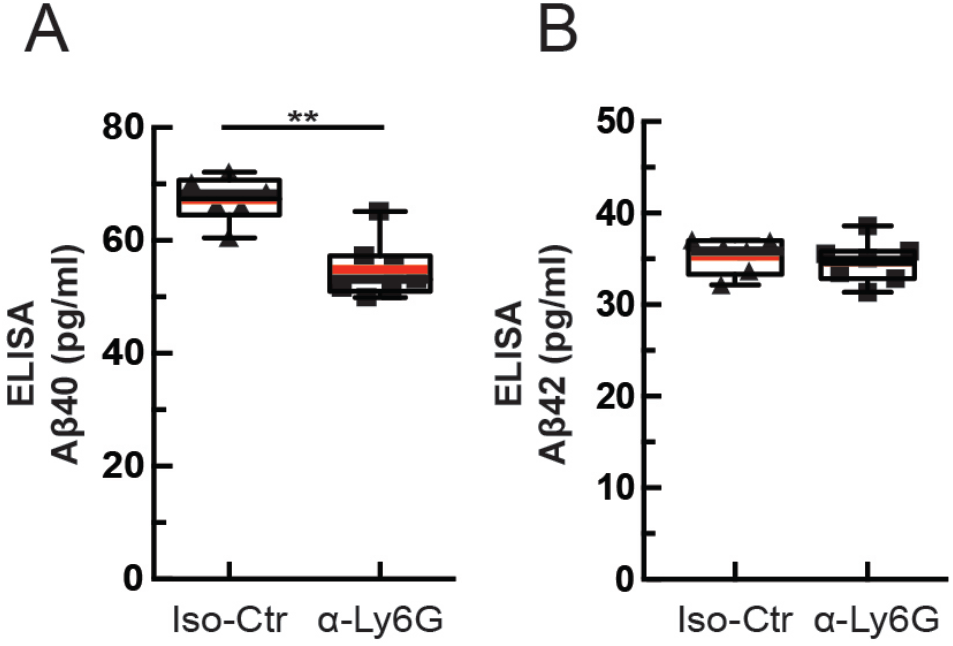
Administration of α-Ly6G for one month decreased the concentration of Aβ1-40 in APP/PS1 mice. ELISA measurements of Aβ1-40 (left) and Aβ1-42 (right) monomer concentrations after 4 weeks of treatment every three days.

Finally, we address the question of how only ~2% of capillaries being stalled could explain the dramatic blood flow changes we observed after anti-Ly6G administration. Because each occluded capillary decreases blood flow in up- and down-stream vessels*^20^*, a small number of stalled capillaries could have an outsized impact on CBF. To estimate the magnitude of this impact and to compare how the topology of the cortical capillary network influences the result, we simulated blood flow in vascular networks from a 1 mm^3^ volume of mouse parietal cortex (Fig. 6a)*^21^*, a 6 mm^3^ volume of human cortex (Fig. 6b)*^22^*, and a synthetic periodic network of order three (Extended Data Fig. 17) using a non-linear model of microvascular blood flow*^23^* (see Supplementary Text). cCBF decreased linearly with an increasing fraction of stalled capillaries, without any threshold effect, across all three networks (Fig. 6c), demonstrating that, on average, each single capillary occlusion has a similar, and cumulative, impact on blood flow. Moreover, the slope of the CBF decrease with increasing capillary stalls was almost identical between the mouse, human, and artificial networks, suggesting that capillary stalling may impact CBF similarly across threedimensional capillary networks with three vessels connected at each node. Quantitatively, these simulations predicted a ~5% (10%) deficit in cCBF due to 2% (4%) of capillaries stalled (relative to the case with no capillary stalls), which is smaller than the increase in CBF we observed with 2PEF and ASL-MRI measurements after anti-Ly6G administration.

**Fig. 6.**
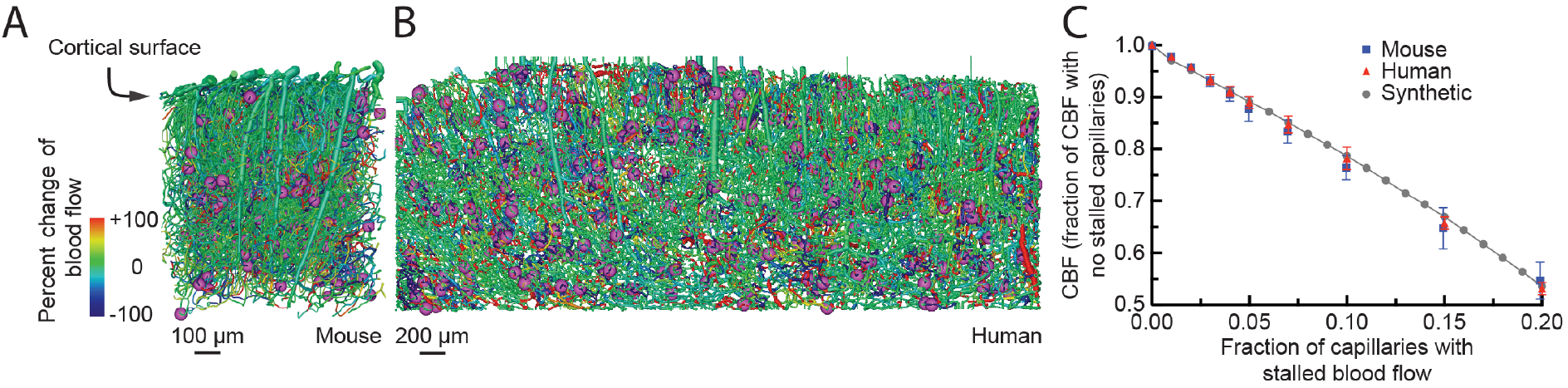
Simulations predicted a similar CBF decrease in mouse and human cortical capillary networks with increasing fraction of capillaries with stalled flow. (A) and (B) Spatial maps of simulated blood flow changes caused by stalling of 2% of capillaries (indicated by purple spheres) in an mouse cortical vascular network (A, from *^40^*), and a human network (B, from *^22^*). (C) Normalized cortical perfusion as a function of the fraction of capillaries that were occluded, expressed as a fraction of the perfusion with no occlusions, in mouse, human, and synthetic networks.

## Discussion

In this study, we aimed to uncover the mechanisms contributing to reduced cCBF in AD and to determine the impact of this reduced cCBF on cognitive function. Brain blood flow reductions occur in the vast majority of dementia patients, including those with AD. These blood flow reductions are one of the earliest features of AD progression*^3,24^*. Mouse models that express mutant APP also show comparable reductions in CBF*^6-8^*.

Previous studies have implicated a variety of potential mechanisms in the CBF reductions seen in AD. Amyloid beta monomers were found to drive vasoconstriction in brain arterioles that could contribute to a reduction in resting CBF^9^. In AD, there is a faster loss of vascular density with age, which could reduce cerebral^10^. In addition to decreases in baseline perfusion, the regulation of blood flow in the brain is compromised in AD. Vessel diameter changes in response to CO_2_ inhalation, blood pressure changes, and changes in local neural activity are all attenuated in AD patients and mouse models of APP overexpression^25^. This loss of dynamic regulation of cerebral blood flow could also contribute to cognitive impacts. Indeed, recent work showed that restoring cerebrovascular function, by angiotensin receptor inhibition or by reducing vascular oxidative stress led to improved cognitive function*^12,26,27^*.

Our data reveal that neutrophil adhesion to the capillary endothelium is a previously unrecognized mechanism for the flow reduction. The rapid resolution of the capillary stalls after α-Ly6G treatment suggests the stalls are caused by receptor-mediated interactions of neutrophils with the capillary endothelium*^28^*, likely due to increased endothelial inflammation. Ly6G has long been appreciated as a neutrophil-specific marker. Consistent with our findings, it has recently been shown that inhibiting Ly6G signaling leads to decreased migration of neutrophils toward sites of inflammation by modulating β2-integrin-dependent adhesion^28^.

Capillary obstructions due to tissue inflammation have been observed in a variety of organ systems (typically at higher incidence than observed here) and have been shown to contribute to the pathology and disease development*^29-35^*. Inflammation is a persistent and well-recognized feature of AD and previous work has demonstrated an increase in inflammatory adhesion receptors on endothelial cells*^36,37^*, which likely underlies the capillary stalling we observed. A significant contributor to this inflammation is increased reactive oxygen species (ROS) induced by brain exposure to Aβ oligomeric aggregates^26^. These ROS cause a loss of cerebrovascular flow regulation and likely drive the expression of leucocyte-binding receptors on the endothelial cell surface, such as ICAM1 and VCAM1. Our observation that some capillary segments were more likely to stall suggests that the underlying vascular inflammation may not be uniform.

While here our focus has been on increased leukocyte adherence causing a subset of capillaries to be transiently stalled due to a firmly adhered leukocyte, this increased leukocyte adherence likely also contributes to slowed, but not stalled, flow in other capillary segments when a leukocyte is present. Our experimental approach does not enable us to readily detect such slowed vessels. Our simulations included only the impact of completely stalled vessels, which may have contributed to the model’s underestimation of the increase in CBF after α-Ly6G administration. However, the simulations predicted a similar sensitivity of brain blood flow to capillary stalling in humans and mice, suggesting that, if capillary stalling occurs in AD patients, significant blood flow improvements could be achieved.

Spatial and working memory improved after increasing blood flow in APP/PS1 mice on a time scale that was too fast for significant neuronal plasticity or synaptic remodeling, suggesting that a mismatch between neuronal energy metabolism and delivery of energy substrates through blood flow contributes to the cognitive deficit. We, and others*^38^*, have also observed improved cognitive function after extended treatment with antibodies that deplete neutrophils. The previous study used different AD mouse models (3xTg and 5xFAD) and different antibodies (α-GR-1, α-LFA-1) than used here and suggested that inhibition of neutrophil trafficking into the brain contributed to the cognitive benefits*^38^*. We have shown that such neutrophil depletion also increases brain blood flow, which may also have contributed to the cognitive improvements in this previous work. Without a firm understanding of the underlying mechanisms that caused reduced CBF in AD, no medical approach to increasing brain blood flow has been developed or tested in humans. In a limited series of experiments in severe AD patients, a piece of omentum, which is known to secrete angiogenic factors and encourage new vessel growth, was surgically placed on the surface of the brain. In the patients that showed an increased CBF as a result, there were signs of improved cognitive function^5,39^. Accordingly, improving CBF by interfering with neutrophil adhesion could be a promising therapeutic approach for AD.

## Acknowledgments

This work was supported by the National Institutes of Health grants AG049952 (CS), NS37853 (CI), and AG031620 (NN), the Alzheimer's Drug Discovery Foundation (CS), the Alzheimer's Art Quilt Initiative (CS), European Research Council grant 615102 (SL), the DFG German Research Foundation (OB), a National Science Foundation Graduate Research Fellowship (JCH), the L’Oréal Fellowship for Women in Science (NN), and used computing resources at CALMIP (SL). We thank Frédéric Lauwers for the human vascular data, Philibert Tsai, Pablo Blinder and David Kleinfeld for the mouse vascular data, and Maria Gulinello for guidance on behavior experiments. Finally, we thank Joseph R. Fetcho, Jesse H. Goldberg, and Michael I. Kotlikoff for commenting on the manuscript. The data reported in this manuscript are archived at **all data underlying these findings will be made publicly available at Cornell’s eCommons online archive**

## Author Contributions

JCCH, OB, SL, NN, and CBS conceived the study. JCCH and CJK performed the *in vivo* imaging experiments. MHJ, GO and YK developed custom software for data analysis. OB conducted the behavioral studies. LP and CI conducted the ALS-MRI experiments. MB, MP, VD, AS, YD and SL performed the blood flow simulations. MCC and SS did the stall analyses in the TgCNRD8 mouse model. JCCH, OB, CJK, VM, LKV, II, YK, JZ, JDB, and ED contributed to the analysis of *in vivo* imaging experiments. JCCH, OB, NN and CBS wrote the paper with contributions from MHJ, MCC, LP, CI, and SL. All authors edited and commented on the manuscript.

## Supplementary Materials for Neutrophil adhesion in brain capillaries contributes to cortical blood flow decreases and impaired memory function in a mouse model of Alzheimer' diseases

### Materials and Methods

#### Animals and surgical preparation

All animal procedures were approved by the Cornell Institutional Animal Care and Use Committee and were performed under the guidance of the Cornell Center for Animal Resources and Education. We used adult transgenic mice (B6.Cg-Tg (APPswe, PSEN1dE9) 85Dbo/J) as a mouse model of AD^41^ and littermate wild-type mice (C57BL/6) as controls (APP/PS1: RRID:MMRRC_034832-JAX, The Jackson Laboratory). Animals were of both sexes and ranged in age from 12 to 100 weeks.

For cranial window implantation, mice were anesthetized under 3% isoflurane on a custom-built stereotactic surgery frame and then maintained on ~1.5% isoflurane in 100% oxygen. Once unresponsive to a toe pinch, mice were given 0.05 mg per 100 g of mouse weight of glycopyrrolate (Baxter Inc.) or 0.005 mg/100 g of atropine (54925-063-10, Med-Pharmex Inc.) to prevent lung secretions, 0.025 mg/100 g of dexamethasone (07-808-8194, Phoenix Pharm Inc.) to reduce post-surgical inflammation, and 0.5 mg/100 g of ketoprofen (Zoetis Inc.) to reduce post-surgical inflammation and provide post-surgical analgesia. Glycopyrrolate and ketoprofen were injected intramuscularly, while atropine and dexamethasone were injected subcutaneously. Bupivacaine (0.1 ml, 0.125%) (Hospira Inc.) was subcutaneously administered at the incision site to provide a local nerve block. Animals were provided 1 ml per 100 g of mouse weight of 5% (w/v) glucose in normal saline subcutaneously every hour during the procedure. We used a thermometer and feedback-controlled heating blanket (40-90-8D DC, FHC) to maintain body temperature at 37 °C. The head was shaved and washed 3 times with alternating 70% (v/v) ethanol and iodine solutions (AgriLabs). A 6-mm diameter craniotomy was performed over the cerebral cortex using a high-speed drill (HP4-917-21, Fordom) using bits with diameters of 1.4, 0.9, 0.7, and 0.5 mm (Fine Science Tools) for different steps in the craniotomy procedure. The craniotomy was then covered with a sterile 8-mm diameter glass coverslip (11986309, Thermo Scientific), glued onto the remaining skull with cyanoacrylate adhesive (Loctite) and dental cement (Co-Oral-Ite Dental). All procedures were done using sterile technique.

Once the craniotomy was completed, mice were returned to their cages and given injections of 0.025 mg/100 g of dexamethasone and 0.5 mg/100 g of ketoprofen subcutaneously 1 and 2 days after surgery, and all cages were placed over a heating pad during this period. Animals were given at least two weeks to recover from cranial window implantation before experimentation to minimize inflammation from the surgical procedure.

Cranial window implantations were also performed in TgCRND8 mice*^17^*. These animals were housed at The Rockefeller University’s Comparative Biosciences Center and treated in accordance with IACUC-approved protocols. The window implantation followed the same protocol as described above, except that mice were anaesthetized using avertin (50 mg/100 g, intraperitoneal) and were given atropine (0.004 mg/100 g).

#### *In vivo* two-photon microscopy

During imaging sessions, mice were anesthetized with 3% isoflurane, placed on a custom stereotactic frame, and were given glycopyrrolate or atropine and glucose as described above. During imaging, anesthesia was maintained with ~1.5% isoflurane in 100% oxygen, with small adjustments to the isoflurane made to maintain the respiratory rate at ~1 Hz. The mouse was kept at 37 °C with a feedback-controlled heating pad (40-90-8D DC, FHC).

To fluorescently label the microvasculature, Texas Red dextran (40 μl, 2.5%, MW = 70,000 kDA, Thermo Fisher Scientific) in saline was injected retro-orbitally just prior to imaging. In some animals, amyloid beta (Aβ) deposits were labeled using methoxy-X04*^42^*. In early experiments using methoxy-X04 obtained directly from Prof. Klunk at the University of Pittsburgh, we retro-orbitally injected 40 μL of 1 mg/ml methoxy-X04 in 0.9% saline (adjusted to pH 12 with 0.1 N NaOH) just prior to imaging. In later experiments using methoxy-X04 available commercially from Tocris, we intraperitoneally injected methoxy-X04 (dissolved in DMSO at 100 mM) one day prior to imaging at a dose of 1 mg/100 g. We observed no obvious differences in the amyloid labeling between these two administration approaches. In some animals, leukocytes and blood platelets were labeled with a retro-orbital injection of Rhodamine 6G (0.1 ml, 1 mg/ml in 0.9% saline, Acros Organics, Pure)*^34^*. Leukocytes were distinguished from blood platelets with a retro-orbital injection of Hoechst 33342 (50 μl, 4.8 mg/ml in 0.9% saline, Thermo Fisher Scientific). Texas Red (and methoxy-X04, when given retro-orbitally) were dosed in a single syringe, while Rhodamine 6G and Hoechst were dosed together in a second syringe.

Three-dimensional images of the cortical vasculature and measurement of red blood cell flow speeds in specific vessels were obtained via a custom-built two-photon excited fluorescence (2PEF) microscope. Imaging was done using 830-nm, 75-fs pulses from a Ti:Sapphire laser oscillator (MIRA HP pumped by a Verdi-V18 or Vision S, Coherent) and 900-nm, 75-fs pulses from a second Ti:Sapphire laser oscillator (Vision S, Coherent). Lasers were scanned by galvonometric scanners (1 frame/s) and focused into the sample using a 20× water-immersion objective lens for high-resolution imaging (numerical aperture (NA) = 1.0, Carl Zeiss Microscopy; or NA = 0.95, Olympus), or a 4× objective for mapping of the cortical surface vasculature (NA = 0.28, Olympus). The emitted fluorescence was detected on either a two-channel detection system or, for later data sets, on an upgraded four-channel detection system. On the two-channel system, the fluorescence was split by a 600nm long pass dichroic and two successive image stacks were acquired first with 645/45 nm (center wavelength/bandwidth) and 575/25 nm bandpass filters to image Texas Red and Rhodamine 6G, respectively, and then with 645/65 nm and 460/50 nm filters to image Texas Red and both methoxy-X04 and Hoescht (on the same channel), all under 830-nm excitation. On the four-channel system, a secondary long-pass dichroic at 520 nm was followed by tertiary long-pass dichroics at 458 nm and one at either 562 or 605 nm. Emission was detected on four photomultiplier tubes through the following emission filters: 417/60 nm for Hoechst, 494/41 nm for methoxy-X04, 550/49 nm for Rhodamine 6G, and 641/75 nm for Texas Red. Laser excitation was 830 nm except when trying to image deep cortical tissue in animals where only Texas Red was present in which case 900-nm excitation was used. Laser scanning and data acquisition was controlled by ScanImage software*^43^*. To visualize the cortical vasculature, stacks of images spaced by 1 μm axially were taken to a cortical depth of 300-500 μm.

For TgCRND8 mice, imaging was performed using a Fluoview 1000MPE two-photon laser scanning microscope (Olympus) equipped with a SpectraPhysics MaiTai DeepSee laser and a 25x/1.05 NA objective at The Rockefeller University BioImaging Resource Center.

#### Awake imaging

A subset of mice was imaged with 2PEF without anesthesia. During the craniotomy surgery, a 3D-printed skull-attached mounting frame was secured on top of the cranial window to allow for head fixation during anesthesia-free imaging. The 3D-printed frame was flanked by 4 screws (TX000-1-1/2 self tapping screws, Small Parts Inc., Miami Lakes, FL) inserted into the skull. The screws and appropriate parts of the frame were glued to the skull using Loctite and dental cement to firmly attach the mounting frame.

We adapted and modified the awake imaging system from Dombeck et al.*^44^*, in which a large (8 inch diameter) Styrofoam ball (Floracraft) was levitated using a thin cushion of air between the ball and a custom made (3D printed) casting containing eight 0.25 inch diameter air jets, arranged symmetrically. The air pressure was adjusted to just float the ball when the mouse was on top of it.

We trained mice to remain in a calm state during awake, head-fixed imaging. During the first training session, mice were handled, with the room lights on, by a trainer wearing gloves for ~10 min or until the mice routinely ran from hand to hand. The mice were then transferred to the ball and allowed to move freely for ~10 min with the room lights on while the handler rotated the ball to keep the mice centered near the top. The second training session consisted of again allowing the mice to move freely on the ball for ~10-15 min, again with the room lights on. The third training session began by head restraining the mice on the ball in complete darkness for ~15-20 min. Typically it would take 5-10 min for the mouse to learn to balance and then begin to walk or run. Mice were then head-fixed and placed on the ball during imaging under the microscope. Awake imaging lasted less than 30 min. Following awake imaging, mice were anesthetized as described above and imaging was repeated over the same cortical area to compare capillary physiology between the awake and anesthetized states.

#### Quantification of capillary network topology and capillary segment stalling

The 2PEF images of vascular networks were manually traced in three-dimensions to create a vectorized skeleton that represents the cortical vasculature using custom-written tracing software. The researchers producing these tracings were blinded to the genotype of the animal and any treatment it had received. Volumes of these image stacks where vessels could not be readily identified and traced were excluded from all analysis. These regions were typically deep and near the edges of the imaged volume, or occasionally directly underneath a large surface vessel. Vessel segments were classified as surface and penetrating arterioles, capillaries, or ascending and surface venules. All vessels smaller than 10 μm in diameter were classed as capillaries. Large surface arterioles were distinguished from large surface venules based on morphology (arterioles were smaller diameter, had smoother walls and less tortuosity, and tended to branch more symmetrically and in Y-shape junctions as compared to venules). Other arterioles or venules were classed by tracing their connectivity to these readily identifiable large vessels.

Each capillary segment in these images was then manually classed as either flowing or stalled based on the motion of RBCs during the entire time each capillary was visible in the 3D image stack. The Texas Red dextran labels the blood plasma, but not the blood cells, so RBCs and other blood cells show up as dark patches in the vessel lumen. The motion of these dark patches indicates flowing blood cells. Each capillary segment was visible in a minimum of ~5 successive frames in the 3D image stack, or for ~5 s (capillaries not oriented parallel to the cortical surface were observed for significantly more frames). We scored a capillary segment as stalled if we did not see motion of the RBCs and other cells in the capillary segment over this observation time. This manual scoring of capillaries as flowing or stalled was performed with the researcher blinded to the genotype and treatment status of the animal. In addition, this scoring was performed using only the image data visible on the Texas Red imaging channel. All animals had at least 800 capillary segments scored as flowing or stalled.

Using the traced vascular network, the topologically shortest path from each flowing or stalled capillary to the nearest penetrating arteriole and ascending venule was calculated using Dijkstra’s algorithm*^45^*.

#### Distinguishing causes of capillary stalls

In some animals, once capillary stalls were identified we used the additional fluorescent labels to determine what was blocking blood flow in the capillary segment. Stalled capillary segments with a cell-shaped object labeled with both Rhodamine 6G and Hoechst present were scored as being caused by a leukocyte. Stalled segments with punctate objects labeled with Rhodamine 6G alone were scored as being caused by platelet aggregates. Stalled capillary segments with only RBCs present were classed as RBC stalls (note that some stalls with leukocytes or platelets present also had RBCs present — these were scored as being caused by the leukoctye or platelet aggregate). A small fraction of capillaries had no moving blood cells in them at all. Because we define flowing vessels by the motion of blood cells, these vessels with no cells present at all were also scored as stalled and classed as “empty,” although it is likely there was plasma flow in these vessel segments.

We assessed if the diameter of flowing and stalled capillaries was different, on average. To reduce the salt and pepper noise in the vascular images, we filtered using a 3D 5 × 5 × 5 pixel Gaussian filter. We then corrected for unevenness in the image intensity by filtering the image (85 × 85 pixel sized mean filter) and subtracting this from the Gaussian filtered image. The resulting image was binarized using Otsu’s method*^46^*. Finally, objects smaller than 1000 voxels were eliminated, where voxels were considered part of the same connected object whenever they shared at least a corner. We then used this binarized image to correct the manual tracing of the vasculature by shifting the centerline so it was equidistant from the vessel boundaries (done within a 10-μm neighborhood to avoid confusion between neighboring capillaries). Every 5 μm along the centerline of each capillary segment, we estimated the vessel radius by finding the closest distance from the centerline to the vessel boundary. Measurements of less than 2 μm or more than 10 μm were excluded as they likely reflected imaging artifacts, and we averaged across all measurements for each capillary segment.

#### Amyloid plaque segmentation and density analysis

2PEF images of methoxy-X04 labeled amyloid plaques were filtered and binarized. Briefly, we first reduced the background signal in a line-by-line fashion by subtracting the median of each line. Salt and pepper image noise was reduced using the adaptive Wiener method with a 3 × 3 pixel kernel*^47^*. The image was then binarized using a manually-determined threshold (99% of the intensity distribution), and smoothed with a 3 × 3 pixel median filter. Objects smaller than 25 voxels were then removed, with object connectivity here defined as voxels sharing a face. The volume fraction of amyloid either globally or in a tube that follows the centerline of each capillary segment was then calculated from this binarized image. The tube volume was generated by swaying a sphere with a specified radius along the centerline of the capillary segment from one end to the other.

#### Kinetics of capillary stalling

To determine the short-term fate of capillaries that stalled, we repeatedly imaged the same capillary bed at baseline and at 5, 10, and 15 min later in APP/PS1 mice (n= 6 animals), and tracked the fate of all the capillaries that were stalled at baseline. If a vessel was observed as stalled at all subsequent imaging time points, it was scored as remaining stalled, and if flow had resumed the stall was scored to have resolved. If the originally stalled capillary resumed flow, then re-stalled at a later time point that was scored as re-stalled. In some animals, we further determined the cause of capillary stalls at each of these time points.

To evaluate the longer-term fate of capillaries that were stalled, we imaged APP/PS1 mice (n= 5 animals) at baseline and then 1, 3, 7, and 14 days later and determined what fraction of the capillaries stalled at baseline were stalled at any subsequent imaging session.

We estimated how frequently we would observe capillaries stalled at baseline to be stalled at any subsequent imaging session assuming that no stalls lasted long enough to stay stalled between imaging sessions and that each capillary segment was equally likely to stall. With this model, the cumulative probability, *P_c_*, of the capillaries stalled at baseline to be stalled at any subsequent imaging session is:

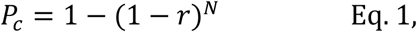

where *r* is the fraction of capillaries with stalled blood flow and *N* is the number of observations after the baseline imaging.

#### Administration of antibodies against Ly6G and impact on neutrophil population

We treated APP/PS1 mice (n = 9, 12-25 weeks old) with intraperitoneal injections of monoclonal antibodies against lymphocyte antigen 6 complex, locus G (Ly6G) (α-Ly6G, clone 1A8, 4 mg/kg, BD Biosciences) or an isotype control antibody (n = 6, Rat IgG2a, κ, 4 mg/kg, BD Biosciences). The same cortical capillary bed was imaged in anesthetized mice immediately before and at 30-60 min and 60-90 min after treatment. Quantification of stalled capillaries was performed blinded to imaging time and treatment type.

To determine the impact of α-Ly6G on neutrophil number, we used flow cytometry to determine neutrophil counts 24 hr after a single treatment (4 mg/kg) and after one month of treatment every three days (2 mg/kg).

Mouse blood was collected from the submandibular vein and mixed with 1x RBC lysis buffer (00-4300-54, ThermoFisher Scientific). After incubation at room temperature for 10 min, the sample was centrifuged at 500 g for 5 min and the supernatant was removed. The cell pellet was re-suspended in 500 uL of Hank’s balanced salt solution (HBSS) supplemented with 1% bovine serum albumin (BSA) and centrifuged again; this washing procedure was repeated 3 times. Following isolation, neutrophils were re-suspended at a density of 10^7^ cells per ml in HBSS supplemented with 1% BSA. The cell samples were labeled at room temperature for 45 min with the following anti-mouse antibodies: anti-CD45 (560695, BD Bioscience), anti-CD11b (557686, BD Bioscience) and anti-Ly6G (551460, BD Bioscience). After washing the samples with HBSS, the remaining leukocytes were analyzed by flow cytometry using a Guava easyCyte Flow Cytometer (EMD Millipore Corporation). Data were analyzed using FlowJo software (FlowJo LLC). The neutrophil population was identified based on the side and forward scatter and later gated for CD45^high^ and CD115^low^ and finally gated as CD11b^high^ and Ly6G^high^ using FlowJo.

#### Measurement of volumetric blood flow in penetrating arterioles

To quantify blood flow in cortical penetrating arterioles, we measured the vessel diameter from image stacks and the centerline RBC flow speed from line-scan measurements, as described in Santisakultarm *T.P.* et al.*^48^*. The volumetric blood flow, *F*, was calculated as:

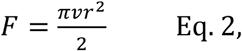

where *v* is the time-averaged centerline RBC speed and *r* is the vessel radius. To correlate the impact of the number of capillaries stalled on penetrating arteriole blood flow, we imaged the same capillaries and measured blood flow in the same six to eight penetrating arterioles in both young APP/PS1 and wt mice (ages 3-4 months) and older APP/PS1 mice (age 11-14 months) treated with antibodies against Ly6G or with isotype control antibodies. Images to determine capillary stalling and line scans to determine penetrating arteriole blood flow speed were taken at baseline and at 30-60 and 60-90 min after treatment. All analysis was conducted blinded to the animal genotype, age, treatment, and imaging time point.

#### Measurement of global blood flow using ASL-MRI

Imaging was performed on a 7.0 Tesla small animal MRI system with 450 mT/m gradient amplitude and a 4500 T/m/s slew rate (Biospec 70/30, Bruker). The animals were anesthetized with isoflurane in oxygen and immobilized in the MRI using a nose cone and bite ring. A volume coil was used for transmission and a surface coil for reception. We imaged APP/PS1 and wt mice (age 7-9 months) at baseline. About 48 hrs later, animals were given an intraperiotenial injection of α-Ly6G or isotype control antibodies (4 mg/kg) and a second set of images were acquired between 2-6 hr after injection.

Anatomical images were acquired to find a coronal slice at a location approximately 1 mm caudal to Bregma*^49^*. This position was used for subsequent ASL imaging, which was based on a FAIR-RARE pulse sequence that labeled the inflowing blood by global inversion of the equilibrium magnetization*^50^*. In this method, inversion recovery data from the imaging slice are acquired after selective inversion of the slice and after inversion of both the slice and the surrounding tissue. The difference of the apparent R1 relaxation rate images then yields a measure of the CBF*^51^*. Three averages of one axial slice were acquired with a field of view of 15 × 15 mm, spatial resolution of 0.23 × 0.23 × 2 mm^3^, echo time TE of 5.36 ms, effective TE of 26.84 ms, repeat time TR of 10 s, and a RARE factor of 36. This resulted in a total scan time for the CBF images of about 25 min. Turbo-RARE anatomical images were acquired with the following parameters: 10 averages of 14 slices with the same field-of-view and orientation as the ASL images, resolution = 0.078 × 0.078 × 1 mm^3^, TE = 48 ms, TR = 2000 ms, and a RARE factor of 10. The total scan time was 6:20 min.

For computation of CBF, the Bruker ASL perfusion processing macro was used. It uses the model and includes steps to mask out the background and ventricles*^52^*. The masked CBF images were exported to Analyze format on the MRI console. We then used the anatomical image to create a mask that outlined the entire cortical region, excluding the sinus, and averaged the CBF measurement across this region for each animal at each imaging time point. Analysis of ASL-MRI data was conducted blinded to animal genotype and treatment.

#### Extraction of network topology and vessel diameters from mouse anatomical dataset

One large postmortem dataset from the vibrissa primary sensory (vS1) cortex in mouse previously obtained by Tsai et al*^21^* and Blinder et al.*^40^*, was used for this study (~1 mm^3^ and ~15,000 vessel segments). In brief, this dataset was obtained by filling the vessels with a fluorescent indicator, extracting the brain and imaging with 2PEF from the pial surface to near the bottom of cortex. In this dataset, penetrating arterioles and ascending venules that reached the pial surface were identified by following their connections to a large cerebral arteriole or venule. We further labeled subsurface vessels in three classes: arterioles, capillaries, and venules. Starting with the surface and penetrating arterioles (venules) were defined by iteratively seeking all vessels with diameter above 6 μm connected to any previously labeled arteriole (venule). All remaining vessels were labeled as capillaries. The diameter threshold was manually chosen as the smallest integer diameter value which resulted in arteriolar and venular trees that exhibited no loops, in contrast to the very looped capillary network.

Due to post-mortem shrinkage, vessel diameters in this mouse dataset were smaller than those measured *in-vivo,* so required rescaling*^21,40,48^*. As blood flow is highly dependent on vessel diameters, two successive corrections were applied. First, a monotonically increasing function, which tends to one at large diameters, was applied to all vessels diameters:

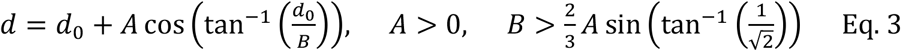

where *d* is the corrected diameter and d_0_ is the diameter extracted from the image stack. *A* and *B* are constrained parameters calculated so that the corrected vessel diameter distribution matched *in vivo* measurements from two photon microscopy^12^, as shown in Extended Data Fig. 18. This function ensures that the hierarchy of diameters in the network is preserved and the larger vessels are not rescaled. For the network represented in Fig. 6a, A=1.4 and B=10.3, so that the diameter threshold for capillary vessels becomes 7.2 μm. A second depth-dependent correction was then applied to the diameter of arterioles and venules:

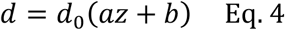

where *z* is the depth below the cortical surface and *a* and *b* are parameters determined so that the diameters of the trunks of the penetrating arterioles and ascending venules matched *in-vivo* measurements^53^. For the network represented in Fig. 6a, these parameters were a=-0.0014 μm^-1^ (-9.36e-4 μm^-1^) and b=2.54 (2.02) for arterioles (venules).

#### Extraction of network topology and vessel diameters from human anatomical dataset

The dataset used was previously obtained by Cassot et al.*^54^* and Lauwers et al.*^22^* from thick sections (300 μm) of a human brain injected with India ink from the Duvernoy collection*^48^*. The brain came from a 60-year old female who died from an abdominal lymphoma with no known vascular or cerebral disease. It corresponds to a large volume (6.4 mm^3^ of cerebral cortex) extending across 20.8 mm^2^ along the lateral part of the collateral sulcus (fisiform gyrus) extracted from Section S2 in Lauwers et al.*^13^*, and includes a total of 27,340 vessel segments. The mean radius and length of each segment were rescaled by a factor of 1.1 to account for the shrinkage of the anatomical preparation. The main vascular trunks were identified manually and divided into arterioles and venules according to their morphological features, following Duvernoy’s classification*^55-56^*. Following Lauwers et al.*^22^* and Lorthois et al*^23^*(*1*) as in the mouse data sets, arterioles (venules) were defined by iteratively seeking all vessels with diameter above 9.9 μm connected to any previously identified arteriole (venule), so that no loops were present. All remaining vessels were classified as capillaries.

#### Synthetic network generation

The synthetic periodic network of order three (i.e. three edges per node) was generated to match the mouse network parameters. A 1-mm^3^ vascular network was constructed by replication of a simple periodic network (Extended Data Fig. 17). Capillary diameters and lengths were uniform and were set to the averages for the mouse network. A single penetrating arteriole and ascending venule (with diameters set to the averages from the mouse network) served as inlet/outlet. The distance between the inlet and outlet corresponded to the average distance between penetrating arterioles and ascending venules from the mouse dataset.

#### Blood flow simulations

The methodology for simulating blood flow in these intra-cortical vascular networks has been presented in detail in Lorthois et al.*^23^*. Briefly, the network was represented by a graph in which edges represent vessel segments between branches that are characterized by an average diameter and length. We used a onedimensional (analogous to electric circuit models) nonlinear network model that was slightly modified from Pries et al.*^57^* to handle large networks for the flow simulations. Using an iterative procedure, the model takes into account the complex rheological properties of blood flow in the microcirculation (Fåhræus, Fåhræus-Lindqvist, and phase separation effects). These effects are modeled using empirical descriptions*^58,59^* deduced from experiments in rats. The model was used to calculate the flow and hematocrit in each vessel and the pressure at each intersection of vessels. For the human dataset, the parameters for the empirical descriptions of the Fåhræus, Fåhræus -Lindqvist and phase separation effects were re-scaled in order to account for the difference in characteristic size between human and rat RBCs, as proposed by Lorthois et al.*^23^* and Roman et al.*^60^*. This simulation approach has no free parameters.

*Boundary conditions:* Physiologically realistic pressure drops of 60 mmHg, as measured in rats*^61^* and estimated in humans*^23^*, were imposed between all arteriolar and venular trunks feeding and draining the computational volume, while a no-flow condition was imposed on deeper arteriolar or venular vessels that intersected the lateral boundaries of the simulated volume. A constant discharge hematocrit of 0.45, corresponding to a typical value of the systemic hematocrit, was also imposed in arteriolar trunks. Moreover, a pseudo-periodic boundary condition was applied to all capillaries at the lateral boundaries, as illustrated in Extended Data Fig. 19. Fictitious vessels were created that link capillaries intersecting opposing faces in a semi random fashion. A grid was created on the two faces and refined until, for a given cell, each capillary on one face was matched with at most 2 capillaries on the opposing face, allowing the creation of fictitious bifurcations. Once the optimal grid was found, the closest neighboring vessels from the opposing faces were connected together. The length of the resulting fictitious vessels was set to 50 μm and their diameters to the average diameters of the connected capillaries. This pseudo-periodic boundary condition is similar in spirit but simpler and more computationally effective than the one recently introduced by Schmid et al.*^62^*. Finally, a no-flow boundary condition was applied to all vessels intersecting the bottom face of the domain. We also compared the results with no-flow boundary conditions for all capillaries at the lateral boundaries.

*Simulating stalls:* In order to study the influence of capillary stalling on cerebral blood flow, a given proportion of capillaries in each network was randomly occluded. To simulate occlusion, the radius of the selected vessels was divided by 100. This resulted in a large increase of the hemodynamic resistance, of order 10^8^, and a similar decrease of the computed flow through these vessels. At least five repetitions were performed for each proportion of stalled capillaries and each set of conditions considered. On the mouse data, 1000 simulations in total were run on a 32-core Intel(R) Xeon E5-2680 v2 @ 3.3 GHz for a total computational time of ~170 hours. For human data set, about 100 simulations were run on the same machine for a total computational time of ~50 hours.

#### Behavior Experiments

All experiments were performed under red light in an isolated room. The position of the mouse’s nose was automatically traced by Viewer III software (Biobserve, Bonn, Germany). In addition to the automatic results obtained by Viewer III software, a blinded experimenter independently scored mouse behavior manually. Animals were taken into the behavior room one-hour prior to the experiment. Behavioral analysis was conducted at baseline and at 3 and 24 h after injection with α-Ly6G or isotype control antibodies (IP 4 mg/kg). Animals were then treated every three days for four weeks (IP 2 mg/kg) and behavior experiments were repeated. The OR, Y-maze, balance beam walk, and forced swim tests were performed at all time points. The NOR task was performed only at baseline and the 4-week time point to avoid animals becoming accustomed to the objects. Animals were ~11 months of age at the start of the experiment (APP/PS1, α-Ly6G n=11; APP/PS1 Iso-Ctl, n=9; wt α-Ly6G, n=10; and wt Iso-Ctl, n=10).

*Object replacement test:* The object replacement (OR) task evaluated spatial memory performance. All objects were validated in a separate cohort of mice to ensure that no intrinsic preference or aversion was observed and animals explored all objects similarly. Exploration time for the objects was defined as any time when there was physical contact with an object (whisking, sniffing, rearing on or touching the object) or when the animal was oriented toward the object and the head was within 2 cm of the object. In trial 1, mice were allowed to explore two identical objects for 10 min in the arena and then returned to their home cage for 60 min. Mice were then returned to the testing arena for 3 min with one object moved to a novel location (trial 2). Care was taken to ensure that the change of placement alters both the intrinsic relationship between objects (e.g. a rotation of the moved object) and the position relative to internal visual cues (e.g. new location in the arena; one wall of testing arena had a pattern). In addition to using the tracking software to determine the object exploration times, the time spent at each object was manually scored by an independent experimenter who was blinded to the genotype and treatment. The preference score (%) for OR tasks was calculated as ([exploration time of the novel object]/[exploration time of both objects]) × 100 from the data in trial 2. Automated tracking and manual scoring yielded similar results across groups, so we report the automated tracking results.

*Y-Maze:* The Y-Maze task was used to measure working memory by quantifying spontaneous alternation between arms of the maze. The Y-maze consisted of three arms at 120° and was made of light grey plastic. Each arm was 6-cm wide and 36-cm long, and had 12.5-cm high walls. The maze was cleaned with 70% ethanol after each mouse. A mouse was placed in the Y-maze and allowed to explore for 6 min. Mouse behavior was monitored, recorded, and analyzed using the Viewer software. A mouse was considered to have entered an arm if the whole body (except for the tail) entered the arm and to have exited if the whole body (except for the tail) exited the arm. If an animal consecutively entered three different arms, it was counted as an alternating trial. Because the maximum number of triads is the total number of arm entries minus 2, the spontaneous alternation score was calculated as (the number of alternating triads)/(the total number of arm entries – 2).

*Forced swim test:* The forced swim test measured depression-like behavior. Mice were individually placed in a 4-L glass beaker filled with 2.5 L of 25°C water. Mice were allowed to adjust for 1 min and then were evaluated for 6 min. An experimenter blind to the genotype and treatment analyzed the videotaped behavior and scored the immobility time, defined by the absence of active, escape-oriented behaviors such as swimming, jumping, rearing, sniffing or diving.

*Balance beam walk:* The balance beam walk measured motor coordination and balance by scoring the ability of the mice to traverse a graded series of narrow beams to reach an enclosed safety platform. The beams consisted of long strips of wood (80 cm) with round cross section of 12- and 6-mm diameter. The beams were placed horizontally, 40 cm above the floor, with one end mounted on a narrow support and the other end attached to an enclosed platform. Bright light illuminated the end of the beam where the mice started. Mice received three consecutive trials on each of the round beams, in each case progressing from the widest to the narrowest beam (15 min between each trial). Mice were allowed up to 60 s to traverse each beam. The time to traverse each beam and the number of times either hind paw slipped off each beam were recorded for each trial. Analysis of each measure was based on the mean score across all trials for that mouse at that time point. Experimenters were blinded to the genotype and the treatment of the mice.

*Novel object recognition test:* The novel object recognition (NOR) task measures recognition memory and is based on rodents’ innate preference for exploring novel objects. This test was conducted only in the animals at baseline and after 4 weeks of treatment. The testing approach was identical to the OR task, but with a novel object placed at the location of one of the initial objects in trial 2.

#### ELISA Assay

After the conclusion of the behavior experiments, the APP/PS1 animals that had received α-Ly6G or isotype control antibodies every 3 days for a month were sacrificed by lethal injection of pentobarbital (5 mg/100 g). Brains were quickly extracted and divided along the centerline. One half was immersed in 4% paraformaldehyde in phosphate buffered saline (PBS) for later histological analysis and the other half was snap frozen in liquid nitrogen.

The frozen APP/PS1 mouse hemi-brains (Iso-Ctr: n=6, 11.5-12.5 months old; α-Ly6G: n=7, 11.5-12.5 months old) were weighed and homogenized in 1 ml PBS containing complete protease inhibitor (Roche Applied Science) and 1 mM AEBSF (Sigma) using a Dounce homogenizer. The homogenates were then sonicated and centrifuged at 14,000 g for 30 min at 4° C. The supernatant (PBS-soluble fraction) was removed and stored at -80° C. The pellet was re-dissolved in 0.5 ml 70% formic acid, sonicated, and centrifuged at 14,000 g for 30 min at 4° C, and the supernatant was removed and neutralized using 1M Tris buffer at pH 11. Protein concentration was measured in the PBS soluble fraction and the formic acid soluble fraction using the Pierce BCA Protein Assay (Thermo Fischer Scientific). The PBS soluble fraction extracts were diluted 1:5. Formic acid extracts were diluted 1:1 after neutralization. These brain extracts were analyzed by sandwich ELISA for Aβ1-40, Aβ1-42, and Aβ aggregates using commercial ELISA kits and following the manufacturer’s protocol (Aβ1-40: KHB3481; Aβ1-42: KHB3441; Aβ aggregates: KHB3491, Thermo Fisher Scientific). The Aβ concentration was calculated by comparing the sample absorbance with the absorbance of known concentrations of synthetic Aβ1-40 and Aβ1-42 standards on the same plate. Data was acquired with a Synergy HT plate reader (Biotek) and analyzed using Gen5 software (BioTek) and Prism (Graphpad).

#### Histopathology

Immunohistochemistry was performed on the brains of mice chronically treated every third day for 4 weeks with either α-Ly6G antibody or isotype control (Iso-Ctr n=5, α-Ly6G n=4). A single paraformaldehyde-fixed hemisphere of each brain was cut into 40 μm thick sagittal sections.

Every sixth section from each mouse was stained with 1% Thioflavin-S (T1892, Sigma) for 10 min at room temperature and washed twice with 80% ethanol for 2 min. The sections were mounted using Fluoroshield with DAPI (F6057, Sigma). Images were taken using confocal microscopy (Zeiss Examiner.D1 AXIO). For each image, the background was subtracted using the ImageJ background subtraction plugin (Rolling ball with 7 μm radius). Images were then manually thresholded, using the same threshold for all sections from a given mouse. Appropriate thresholds varied mouse to mouse and were set to ensure that the smallest Thioflavin-S labeled objects that morphologically appeared to be an amyloid plaque remained above threshold. Cortical and hippocampal regions of interest were defined in each section anatomically, and the fraction of pixels above threshold was determined across all sections for these regions of interest. All image processing was done blinded to treatment group. As a second measure of amyloid deposition, we manually counting the number of Thioflavin-S positive amyloid plaques in the cortex and hippocampus, again across all sections and while blinded to the treatment group. All sections were stained and imaged in parallel. Artifacts such as bubbles were eliminated from analysis by manually excluding these regions.

#### Statistical analysis

Boxplots were created using Prism7 (GraphPad). The box extends between the values for the 25th and 75th percentile of the data. The whiskers extend 1.5 times the difference between the value of the 75th and 25th percentile of the data from the top and bottom of the box. Values lying outside the whiskers were defined as outliers and the mean was computed excluding these outliers. The median is indicated with a black horizontal line inside the box, while the mean is indicated with a red horizontal line. Violin plots were created using the statistical software package, R*^63^*.

Data in all groups was tested for normality using D’Agostino-Pearson omnibus normality test. Parametric statistics were used only if the data in all groups in the comparison were normally distributed. The statistical significance of differences between multiple groups was determined using one-way analysis of variance (ANOVA) followed by Tukey’s multiple comparison correction for normally distributed data, and using Kruskal-Wallis one-way ANOVA followed by Dunn’s multiple comparison correction for data with a non-normal distribution. Statistical comparisons between two groups were performed using the Student’s t test or paired t test for normally distributed data, or using the Mann-Whitney test or Wilcoxon matched-pairs test for data with a non-normal distribution. P-values smaller than 0.05 were considered statistically significant. All statistical analysis was performed using Prism7 (GraphPad).

We use a standardized set of significance indicators across all figures in this manuscript. For comparisons between groups: *p<0.05, **p<0.01, ***p<0.001, ****p<0.0001. For matched comparisons before and after treatment: +p<0.05, ++p<0.01. Supplementary Table 1 provides details of the groups compared, animal and capillary numbers, statistical tests, and explanatory notes for individual panels in the main figures. This information is included in the caption of supplementary figures.

### Supplementary text on numerical simulations of cerebral blood flow changes induced by capillary occlusions

In previous work, we studied how the occlusion of a single cortical capillary influenced blood flow in downstream vessels^11^ and found strong reductions in blood flow (10% of baseline value 1 branch downstream; 25% at 2 branches; 50% at 3 and 4 branches), suggesting that even the small fraction of occluded capillaries we observed in APP/PS1 mice could cause a significant decrease in overall brain blood flow. To test this idea, we simulated blood flow in anatomically accurate blood vessel networks from mice and humans and examined how flow changed when we occluded a random selection of capillaries.

*Validation of simulations by comparison to in vivo measurements in mouse:* As described in the Materials and Methods above, our simulations resulted in calculated values for flow (Extended Data Fig. 20a), pressure (Extended Data Fig. 20b), and hematocrit (Extended Data Fig. 20c) in each vessel segment in the volume. We validated the simulation by comparing *in vivo* measurements of blood flow at different levels in the microvascular hierarchy acquired by 2PEF from the top 300 μm of mouse cortex (data from Santisakultarm, *et al.^48^*) with the simulation predictions. The simulation results are highly dependent on the boundary conditions imposed on capillaries at the lateral edges of the simulation volume. The calculated velocity distribution using pseudo-periodic boundary conditions in capillaries up to 300 μm in depth and using the vessel diameter corrections described above matches the experimental distribution well (Extended Data Fig. 20d). For comparison, the velocity distribution calculated using diameters from the raw datasets (without correction for the difference in vessel size between *in vivo* and post mortem measurements) and that calculated using a no-flow boundary condition both led to an order of magnitude underestimation of capillary flow speeds (Extended Data Fig. 20d). Our new pseudo-periodic boundary condition, together with the correction of vessel diameters, led to a velocity distribution that approaches the distribution of experimental velocities. The experimental distribution has a sharper peak, which might be due to experimental bias associated with the limited number of vessels in which these measurements have been performed (147 *in vivo* measurements vs. 3,400 capillaries in the simulations). The simulated speeds in penetrating arterioles and ascending venules as a function of their diameters also closely matched experimental results from Santisakultarm, et al.*^48^* and from Taylor, et al.*^53^* (Extended Data Fig. 20e).

*Numerical simulation of cerebral blood flow reductions caused by capillary occlusions:* The effect of occlusions in capillaries was investigated by randomly selecting a given proportion of capillaries and reducing their flow by imposing a 100-fold reduction in diameter (Extended Data Fig. 21a, Fig. 6a). To quantify the effects of the occlusions, we calculated the normalized cortical perfusion as the summed flow in the penetrating arterioles feeding the region, normalized by the value calculated with no capillary occlusions (Fig. 6c). While the magnitude of this summed flow is highly dependent on the boundary conditions, the decrease in flow due to capillary occlusions was much less sensitive to the choice of boundary conditions (Extended Data Fig. 21B). For the mouse network shown in Figs. 6a and Extended Data 20a with pseudo-periodic boundary conditions and diameter correction, we found a linear decrease in the normalized perfusion with a slope S=-2.3±0.2 %baseline perfusion/% capillaries stalled (mean±SD) (Fig. 6c). This linear behavior was very robust to variations in the parameters chosen for the computations, with slopes equal to -2.2±0.1 (-2.1±0.2) with no-flow boundary conditions and diameter correction (no diameter correction). In order to evaluate the influence of boundary conditions with regard to the size of the simulated volume, 300 μm-thick subvolumes of the mouse anatomical datasets were randomly extracted. The decrease in blood flow with increasing numbers of stalled capillaries was slightly larger when 300 μm-thick sub-volumes of the datasets were used (-2.6±0.4 and -2.9±0.5 with the pseudo-periodic boundary condition and the no-flow boundary condition, respectively), as compared to the full ~1 mm-thick volume. In Fig. 1j, only computations on the maximum simulation volume with the corrected diameters and pseudo-periodic boundary conditions are presented.

The simulations in the human network (Fig. 6b) using pseudo-periodic boundary conditions yielded a slope of S=-2.3±0.6, very similar to the mouse results. This linear decrease was also observed in synthetic periodic networks of order three (i.e. three edges per node; S=-2.9, Fig. 6c).

*Limitations and methodological considerations:* The human dataset used in the simulations was only 300 μm thick, raising concerns about the influence of boundary conditions. The broad agreement between simulation results in mouse datasets with 1-mm and 300-μm thickness reduces this concern. The simulations predicted a similar CBF increase across mouse and human vascular networks when stalls were reduced, suggesting that the blood flow improvements we observed in APP/PS1 mice may be achievable in humans.

The simulations predicted a smaller impact of capillary stalling on CBF than we observed experimentally. One possible explanation is that the simulations used vascular networks from wt mice, while AD mouse models have different vascular densities and topologies*^64^* that may influence the sensitivity of CBF to capillary stalls. In addition, increased leukocyte adhesion in APP/PS1 mice may lead not only to complete stalls, but also to slowed flow in some capillaries that is not captured in the simulations.

We also observed differences in the 2PEF and ASL-MRI measurements of blood flow increase after alpha-Ly6G administration that may be due to the limited sampling of penetrating arterioles at one anatomical location with 2PEF and to taking ASL-MRI data several hours later after treatment than 2PEF. In addition, the correlation between ASL-MRI values and arteriole flow may not be exact due to technical limitations in mouse ASL-MRI.

### Supplementary Figures

**Extended Data Fig. 1.**
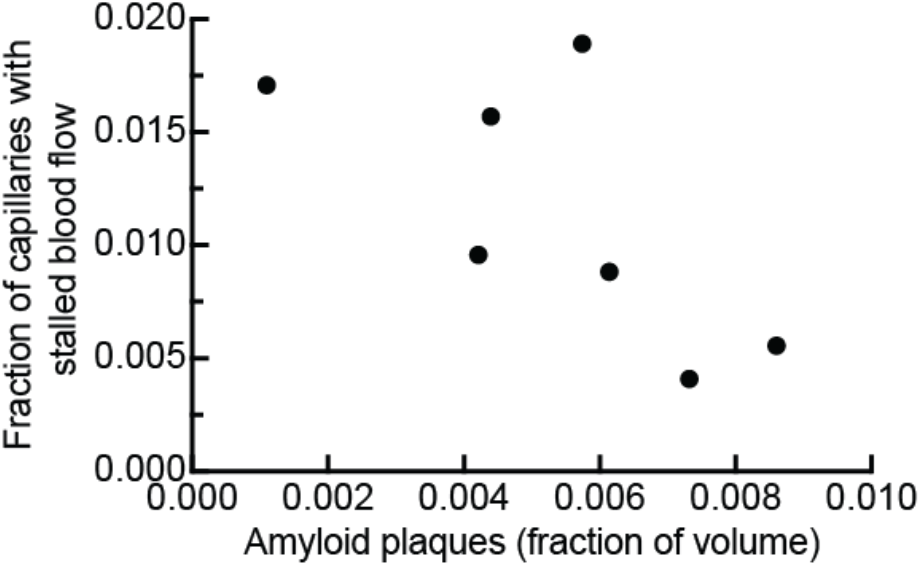
The fraction of capillaries with stalled blood flow did not increase with increasing cortical amyloid plaque density in APP/PS1 mice. Fraction of capillaries with stalled blood flow as a function of the cortical volume fraction that was labeled by methoxy-X04. Mice ranged from 50 to 64 weeks of age.

**Extended Data Fig. 2.**
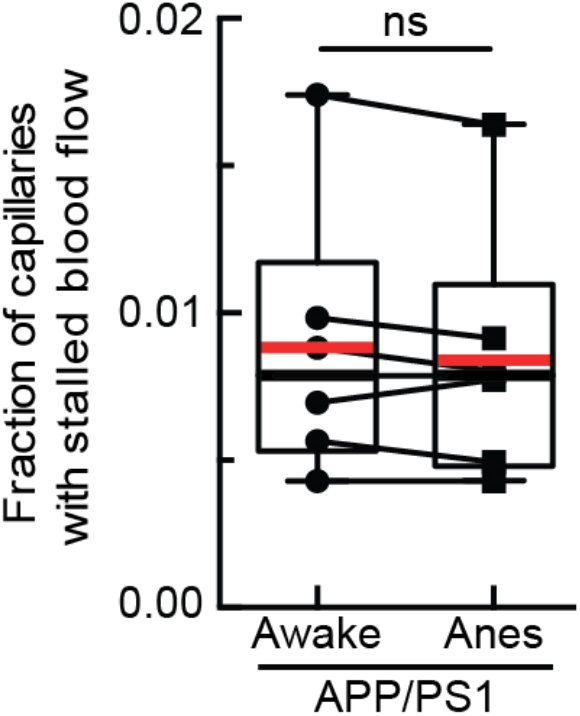
Plot of the fraction of capillaries with stalled blood flow in mice imaged while anesthetized and awake. Lines connecting data points indicate data from the same animal. Animals were first trained to remain calm while head fixed and standing on a spherical treadmill. On the day of imaging, animals were briefly anesthetized to enable retro-orbital injection of Texas-Red dextran, and were then allowed to wake up. We imaged these animals first while awake and then while anesthetized under 1.5% isoflurane, with both imaging sessions occurring on the same day (n = 6 mice, no significant difference by Wilcoxon test).

**Extended Data Fig. 3.**
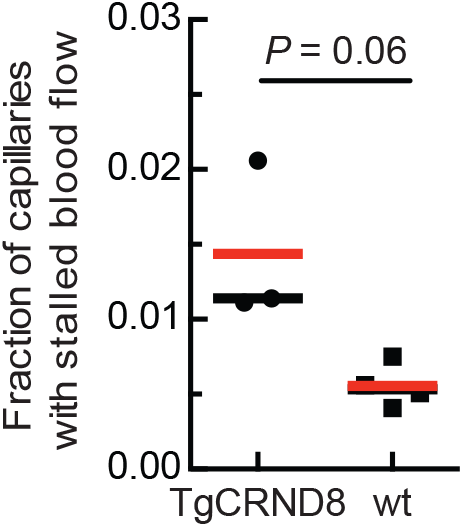
2PEF imaging of cortical vasculature reveals a higher fraction of stalled capillaries in TgCRND8 mice as compared to wt mice. Fraction of capillaries with stalled blood flow in TgCRND8 and age-matched wild type littermates (TgCRND8: 3 mice, 3,028 capillaries; wild type: 4 mice, ~4,062 capillaries; p=0.06, Mann-Whitney).

**Extended Data Fig. 4.**
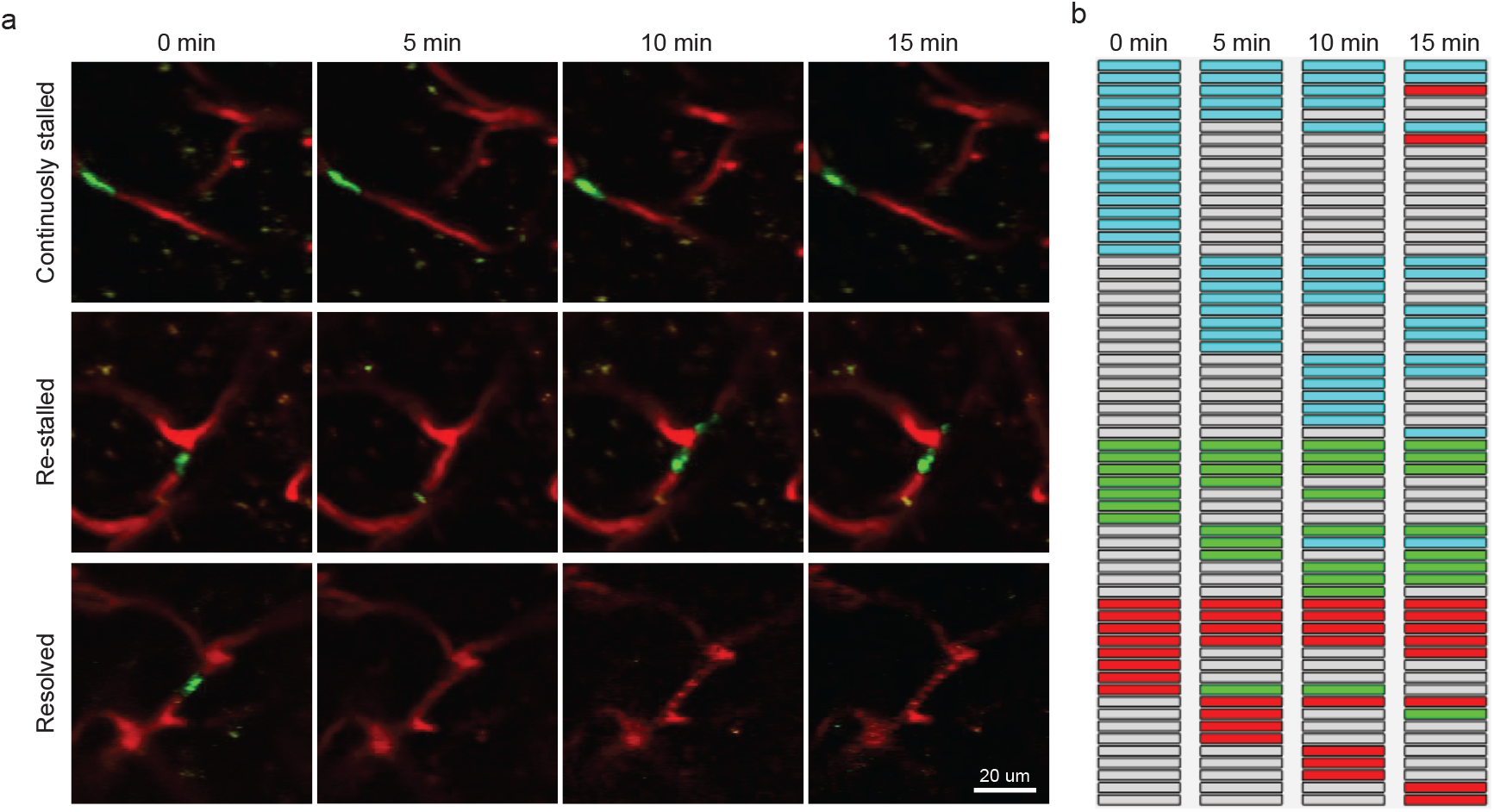
Characterization of capillary stall dynamics in APP/PS1 mice. (a) Repeated 2PEF imaging over 15 min of capillaries that were stalled at the baseline measurement and (top) remained stalled, (middle) began flowing and then re-stalled and (bottom) resolved and remained flowing. Blood plasma labeled with Texas-Red dextran (red) and leukocytes labeled with Rhodamine 6G (green). (b) Characterization of the fate of individual capillaries observed as being stalled across four image stacks taken at baseline and 5, 10, and 15 min later. Each row represents an individual capillary and the color of the box for each capillary at each time point indicates the status: flowing (grey), stalled with a leukocyte present (cyan), stalled with platelet aggregates present (green), and stalled with only RBCs (red).

**Extended Data Fig. 5.**
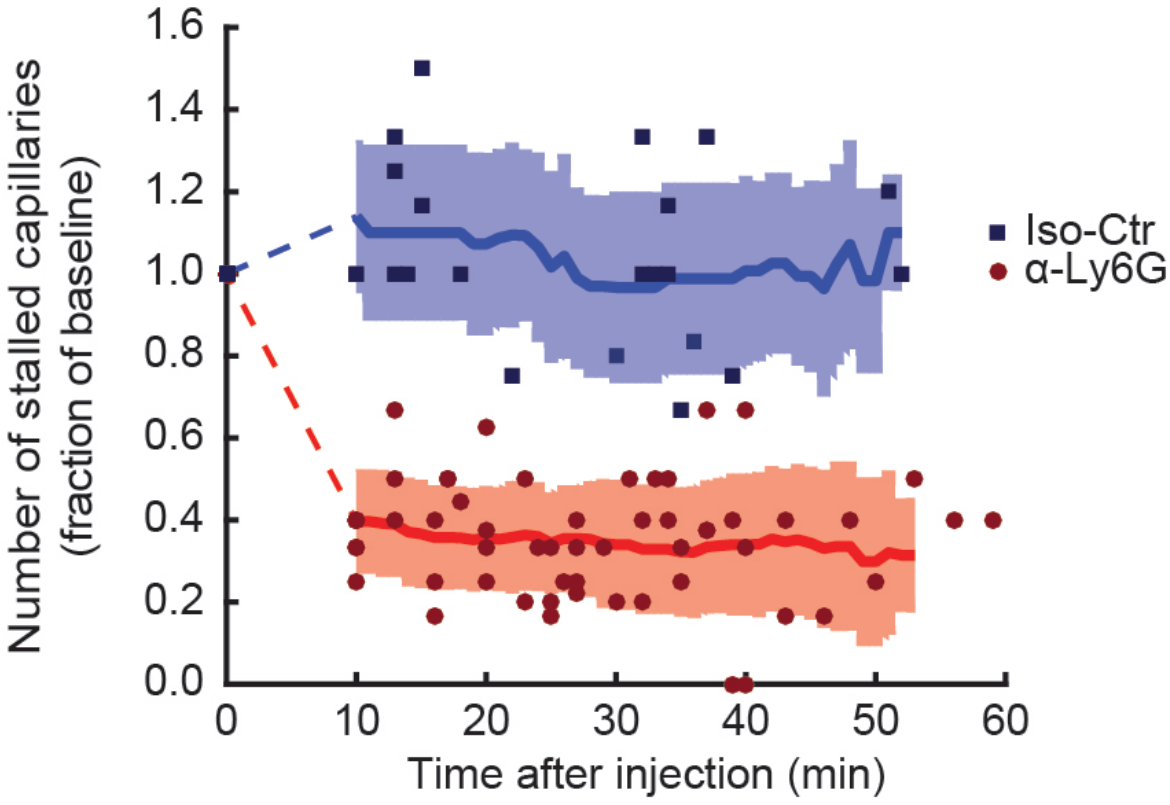
Number of stalled capillaries in APP/PS1 mice dropped rapidly after α-Ly6G administration. 2PEF image stacks were taken repeatedly over an hour after α-Ly6G or isotype control antibody injection and the number of stalled capillaries determined at each time point (α-Ly6G: n=6 mice; Iso-Ctr: n=4; each mouse imaged 2 to 6 times over the hour).

**Extended Data Fig. 6.**
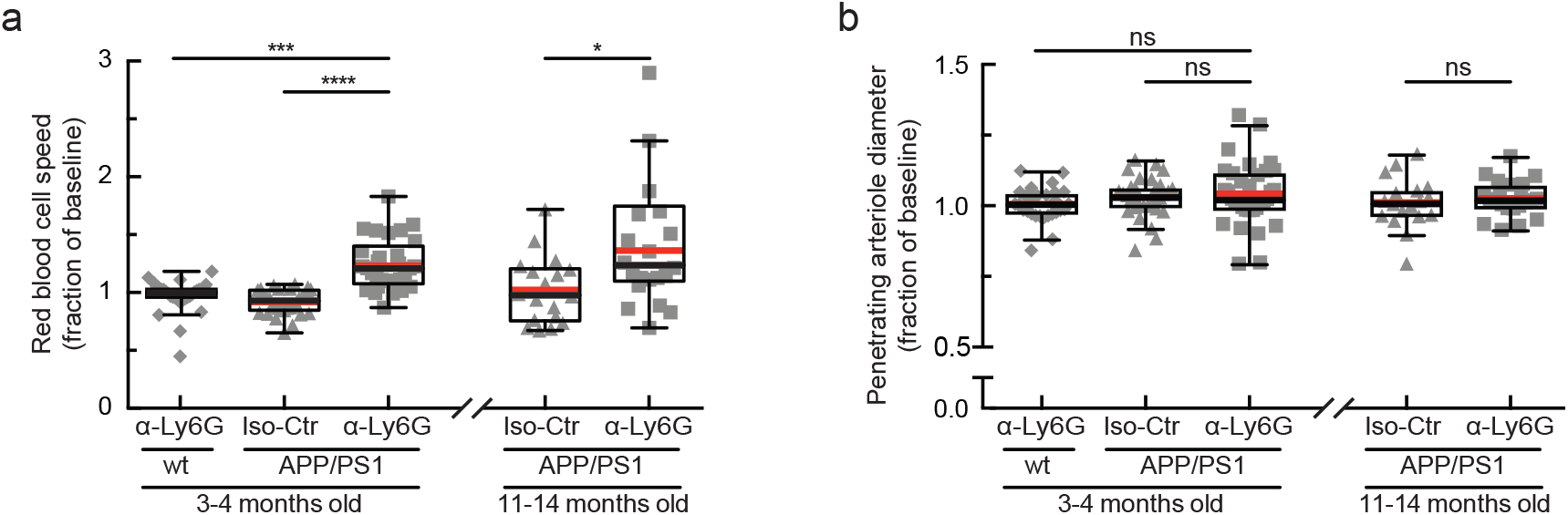
Administration of antibodies against Ly6G increased the RBC flow speed but did not alter the diameter of cortical penetrating arterioles in APP/PS1 mice. (a) RBC flow speed and (b) vessel diameter after α-Ly6G or isotype control antibody administration in young (3-4 months) and old (11-14 months) APP/PS1 mice and wt control animals shown as a fraction of baseline (young wt α-Ly6G: 5 mice, 30 arterioles; young APP/PS1 Iso-Ctr: 5 mice, 32 arterioles; young APP/PS1 α-Ly6G: 5 mice, 33 arterioles; old APP/PS1 Iso-Ctr: 3 mice, 18 arterioles; old APP/PS1 α-Ly6G: 3 mice, 22 arterioles; * p<0.05, ** p<0.01, *** p<0.001, **** p<0.0001, Kruskal-Wallis one-way ANOVA with post-hoc pair-wise comparisons using Dunn’s multiple comparison test).

**Extended Data Fig. 7.**
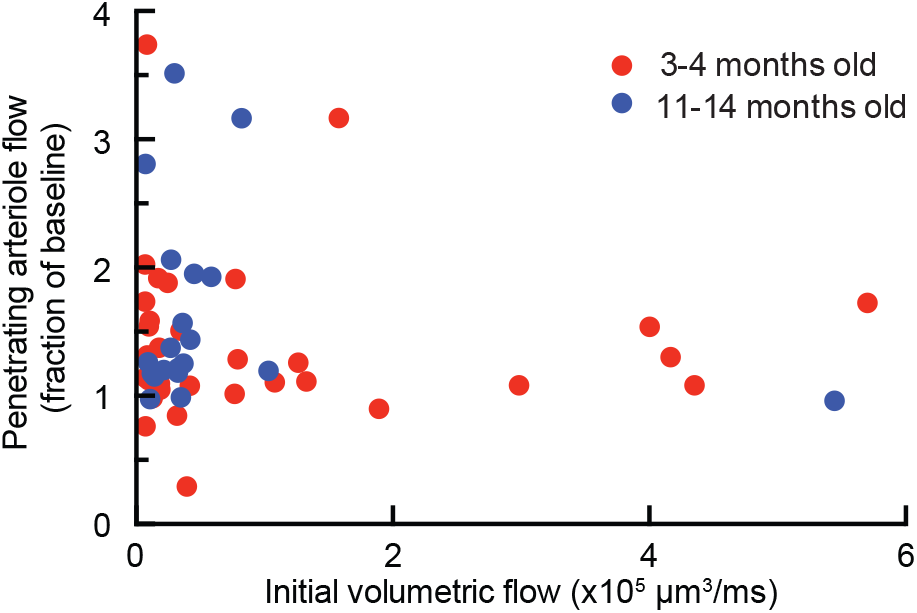
Penetrating arterioles with slower initial flow tended to increase flow speed more after α-Ly6G injection in APP/PS1 mice. Plot of penetrating arteriole flow after α-Ly6G antibody administration in young (3-4 months) and old (11-14 months) APP/PS1 mice shown as a fraction of baseline flow. Same data as shown in Fig. 3C.

**Extended Data Fig. 8.**
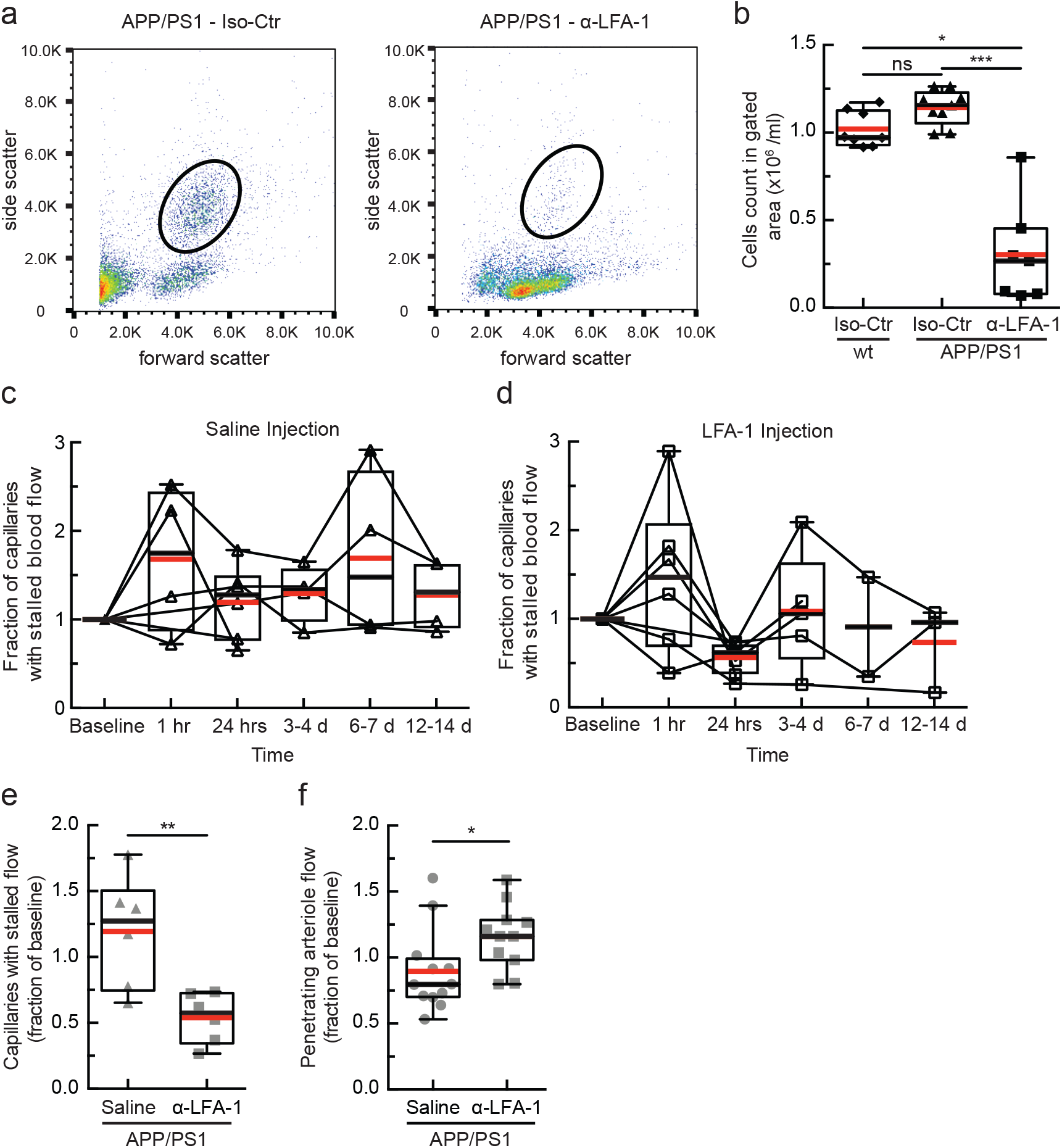
Treating APP/PS1 mice with α-LFA-1 reduced the number of stalled capillaries and improved arterial blood flow after 24 hours. (a) Flow cytometry scatter plots for APP/PS1 mice 24 hours after injection of isotype control antibodies (left) or with antibodies against Lymphocyte Functional Antigen 1 (α-LFA-1; M17/4 clone, BD Biosciences; 4 mg/kg, retro-orbital injection). Circles depict the gate used to identify leukocytes. (b) Leukocyte concentration in the blood 24 hours after treatment with α-LFA-1 or isotype control antibodies in APP/PS1 and wt mice. Leukocytes counts in the gating area were decreased by 84% after α-LFA-1 as compared to the isotype control in APP/PS1 mice (Iso-Ctr in wt: 8 mice, Iso-Ctr in APP/PS1: 9 mice, α-LFA-1 in APP/PS1: 7 mice; *p<0.05, ***p<0.001, Kruskal-Wallis one-way ANOVA with post-hoc pair-wise comparisons using Dunn’s multiple comparison test). (c and c) Fraction of capillaries with stalled blood flow as a function of time after a single retro-orbital treatment with 0.9% saline (c) or α-LFA-1 antibodies (d) in APP/PS1 mice (saline: n = 6 mice; α-LFA-1: n = 7 mice, 4 mg/kg). We observed a transient increase in the number of capillaries with stalled blood flow at about 1 hr after treatment in both groups. There was a significant decrease in the fraction of stalled capillaries 24 hours after injection in the α-LFA-1 group. Images were collected over the same capillary bed on each imaging day, and the fraction of capillaries stalled was determined for each time point, with the analysis performed blinded to treatment day and treatment type. (e) Number of stalled capillaries, expressed as a fraction of the baseline number, 24 hrs after administration of α-LFA-1 or saline. α-LFA-1 reduced capillary stalls by 65% as compared to the saline control. (n = 6 mice per treatment group. **p<0.01, Mann-Whitney test). (f) Fraction of baseline arteriole flow in penetrating arterioles from APP/PS1 mice 24 hours after α-LFA-1 or saline treatment. Each point represents a single arteriole in one mouse. The blood flow was increased after α-LFA-1 treatment by 29% compared with saline controls (APP/PS1 α-LFA1: 4 mice, 11 arterioles; APP/PS1 saline: 4 mice, 12 arterioles; *p<0.05, Mann-Whitney test).

**Extended Data Fig. 9.**
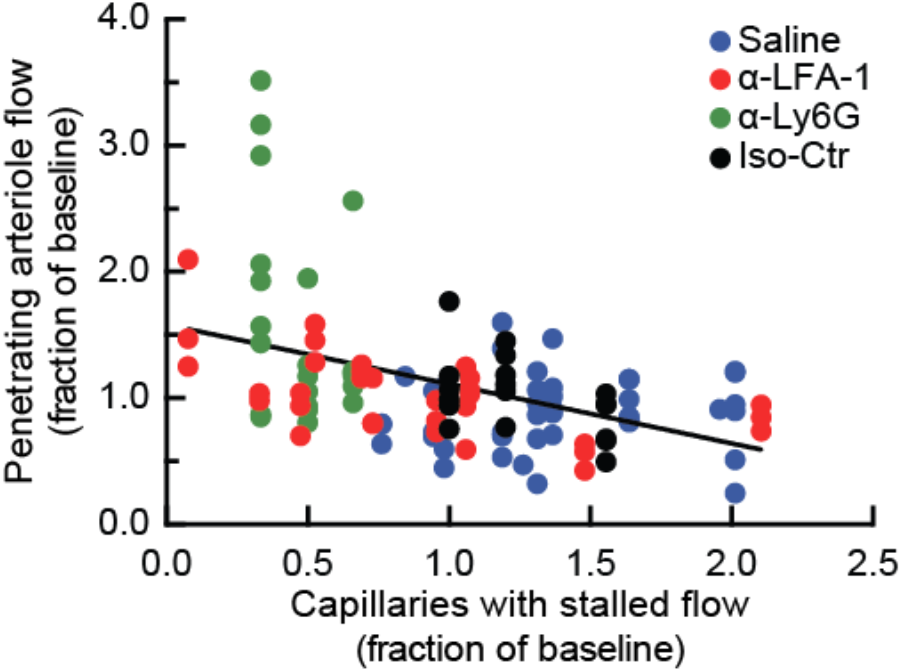
Brain penetrating arteriole blood flow negatively correlates with the number of capillaries stalled in underlying capillary beds in APP/PS1 mice. To correlate the effect of capillary stalling on penetrating arteriole blood flow, we imaged the same capillaries and measured blood flow in the same penetrating arterioles in APP/PS1 mice multiple times before and after administration of saline, α-LFA-1, α-Ly6G, and isotype control antibodies. For saline and α-LFA-1 animals, there were measurements at multiple time points over two weeks (data in Extended Data Fig. 9). For α-Ly6G and isotype control animals there were measurements only at baseline and ~1 hr after administration (data in Fig. 3c and Extended Data Figs. 7 and 8). For each penetrating arteriole at each imaged time point, we plotted the volumetric flow, expressed as a fraction of the baseline volumetric flow, as a function of the number of capillaries stalled at that time point, expressed as a fraction of the baseline number of capillaries stalled (APP/PS1 α-LFA1: 4 mice, 11 arterioles; APP/PS1 saline: 4 mice, 12 arterioles; APP/PS1 α-Ly6G: 3 mice, 22 arterioles; APP/PS1 Iso-Ctr: 3 mice, 18 arterioles). These data confirm the sensitive dependence of penetrating arteriole blood flow on the fraction of capillaries with stalled flow across several different manipulations that led to either increases or decreases in the fraction of capillaries that are stalled. The linear regression is defined by: Y = -0.47 X + 1.6 (R^2^ = 0.2, goodness of fit test; 95% confidence interval on slope: -0.65 – -0.29).

**Extended Data Fig. 10.**
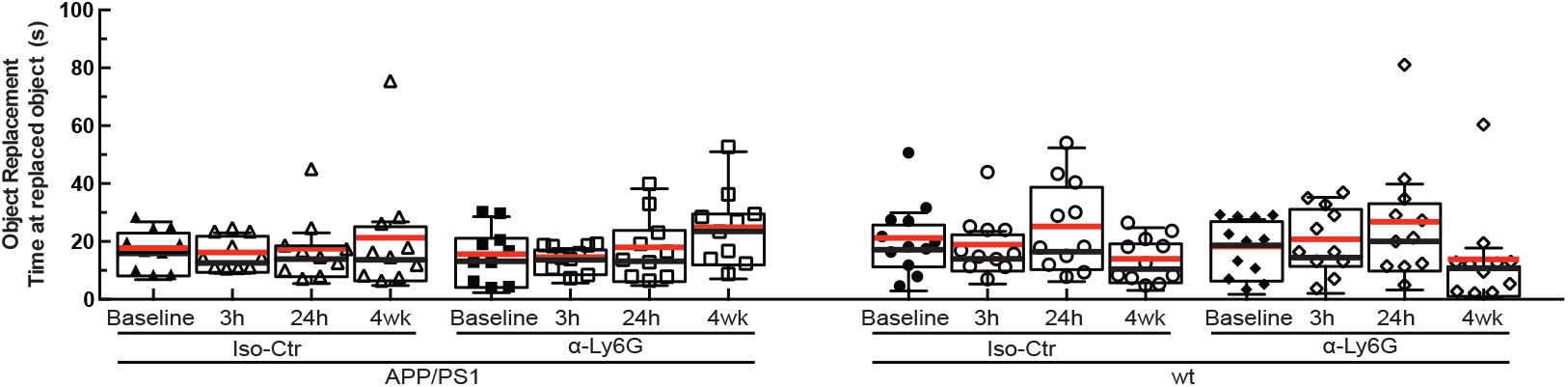
Time spent at the replaced object in wild type controls and APP/PS1 animals treated with α-Ly6G or isotype control antibodies. Time spent at the replaced object measured over 6 minutes for APP/PS1 and wt mice at baseline and at 3h and 24h after a single administration of α-Ly6G or isotype control antibodies, and after 4 weeks of treatment every three days (APP/PS1 Iso-Ctr: 10 mice; APP/PS1 α-Ly6G: 10 mice; wt Iso-Ctr: 11 mice; wt α-Ly6G: 11 mice; no significant differences among groups as determined by Kruskal-Wallis one-way ANOVA).

**Extended Data Fig. 11.**
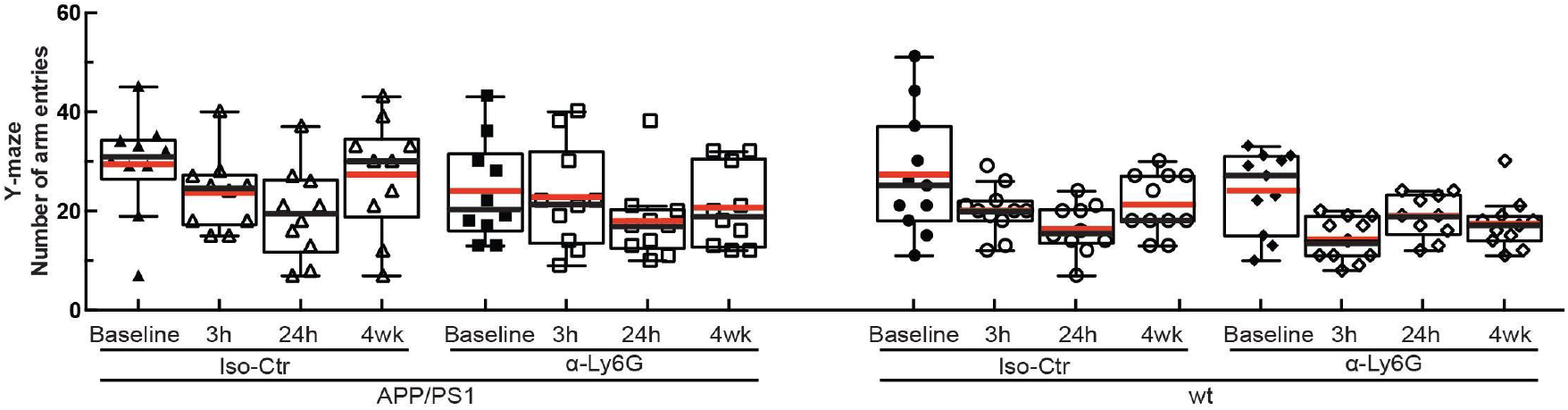
Number of arm entries in the Y-maze for wild type controls and APP/PS1 animals treated with α-Ly6G or isotype control antibodies. Number of arm entries in the Y-maze measured for 6 minutes for APP/PS1 and wt mice at baseline and at 3h and 24h after a single administration of α-Ly6G or isotype control antibodies, and after 4 weeks of treatment every three days (APP/PS1 Iso-Ctr: 10 mice; APP/PS1 α-Ly6G: 10 mice; wt Iso-Ctr: 11 mice; wt α-Ly6G: 11 mice; no significant differences among groups as determined by Kruskal-Wallis one-way ANOVA).

**Extended Data Fig. 12.**
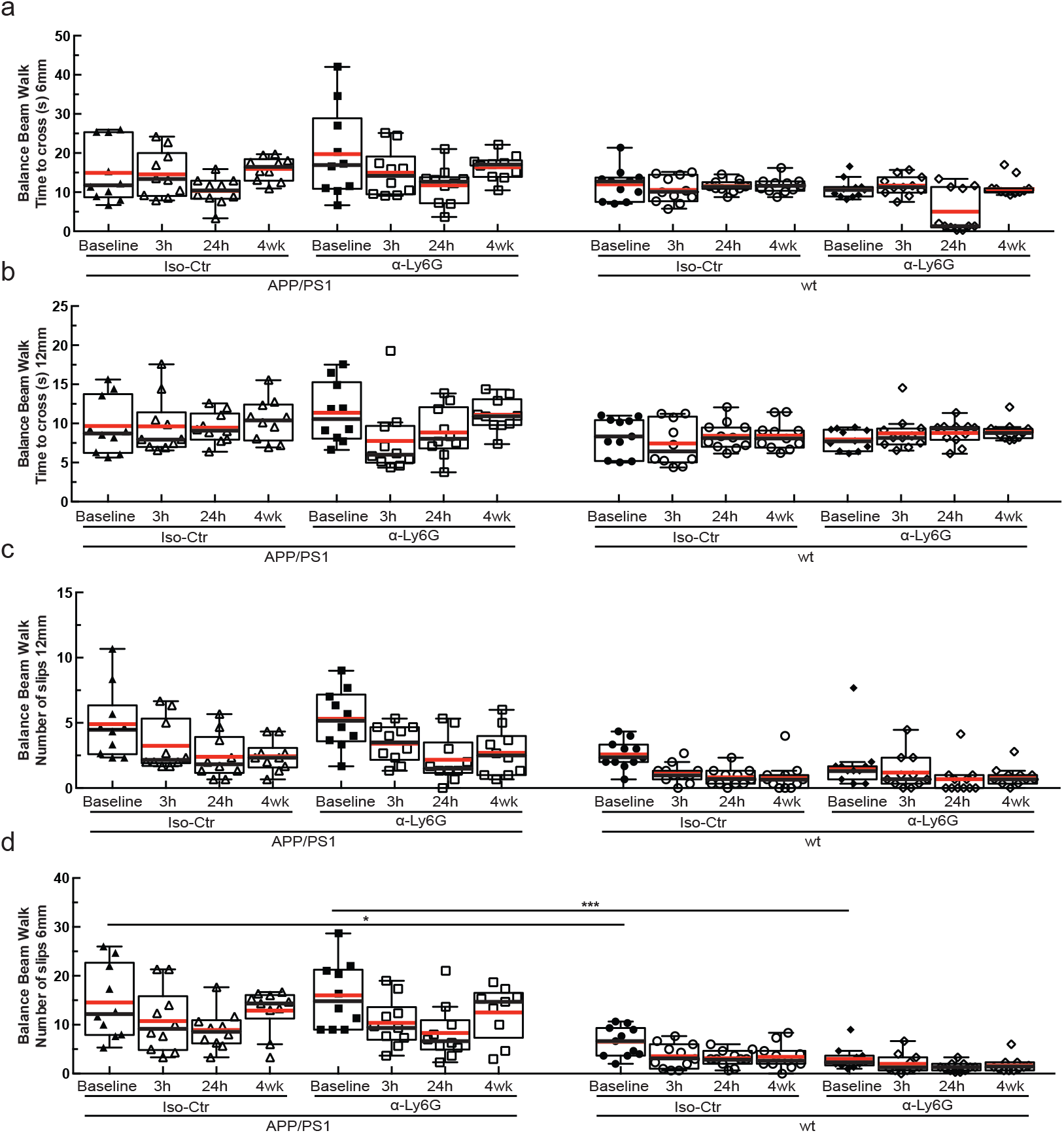
Balance beam walk (BBW) to measure motor coordination in wildtype controls and APP/PS1 animals treated with α-Ly6G or isotype control antibodies. (a and b) BBW time to cross on a 6- and 12-mm diameter beam, respectively, for APP/PS1 and wild type mice at baseline and at 3h and 24h after a single administration of α-Ly6G or isotype control antibodies, and after 4 weeks of treatment every three days. APP/PS1 mice showed a modest trend toward taking more time to cross the 6-mm diameter beam as compared to wt controls. (c and d). Number of slips on the BBW for a 6- and 12-mm diameter beam, respectively, for APP/PS1 and wild type mice at baseline and at 3h and 24h after a single administration of α-Ly6G or isotype control antibodies, and after 4 weeks of treatment every three days. For both beam diameters, APP/PS1 mice showed significantly more slips while crossing the beam as compared to wt animals, suggesting a motor deficit in the APP/PS1 mice. All animal groups showed a reduction in the number of slips with subsequent trials, suggesting improved motor coordination with practice. This improvement did not appear different between α-Ly6G and isotype control treated APP/PS1 mice, suggesting that increases in brain blood flow did not influence the motor learning underlying the reduction in the number of slips (APP/PS1 Iso-Ctr: 10 mice; APP/PS1 α-Ly6G: 10 mice; wt Iso-Ctr: 11 mice; wt α-Ly6G: 11 mice; * p < 0.05, *** p < 0.001, Kruskal-Wallis one-way ANOVA with post-hoc pair-wise comparisons using Dunn’s multiple comparison test)

**Extended Data Fig. 13.**
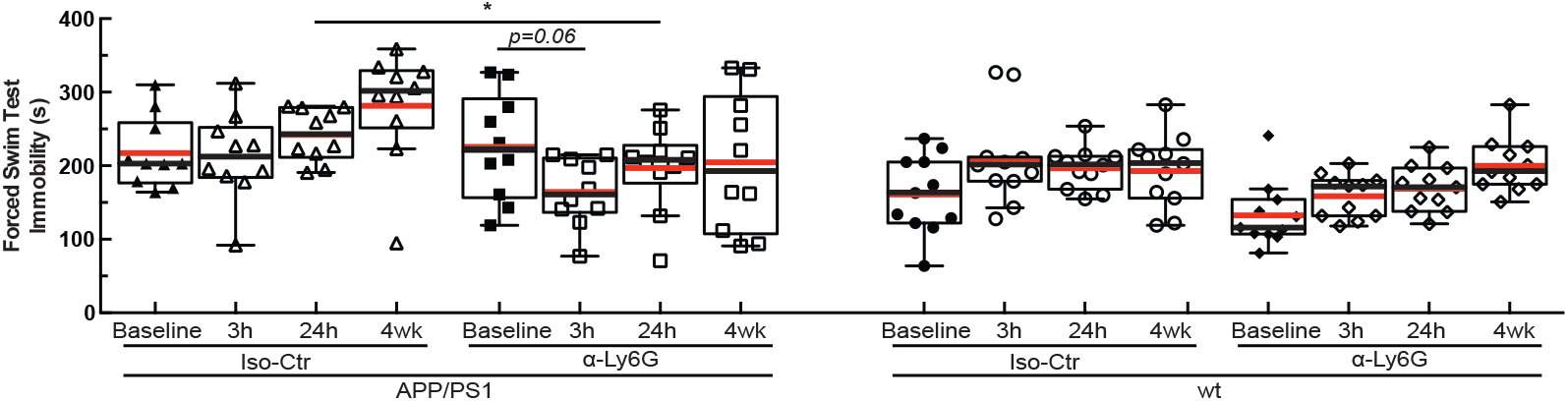
Depression-like behavior measured as immobility time in a forced swim test for wild type controls and APP/PS1 animals treated with α-Ly6G or isotype control antibodies. Immobility time in forced swim test measured over 6 minutes for APP/PS1 and wt mice at baseline and at 3h and 24h after a single administration of α-Ly6G or isotype control antibodies, and after 4 weeks of treatment every three days (APP/PS1 Iso-Ctr: 10 mice; APP/PS1 α-Ly6G: 10 mice; wt Iso-Ctr: 11 mice; wt α-Ly6G: 11 mice; * p < 0.05, Kruskal-Wallis one-way ANOVA with post-hoc pair-wise comparisons using Dunn’s multiple comparison test; p=0.06 comparison between baseline and 3h for APP/PS1 α-Ly6G, Wilcoxon matched-pairs signed rank test relative to baseline).

**Extended Data Fig. 14.**
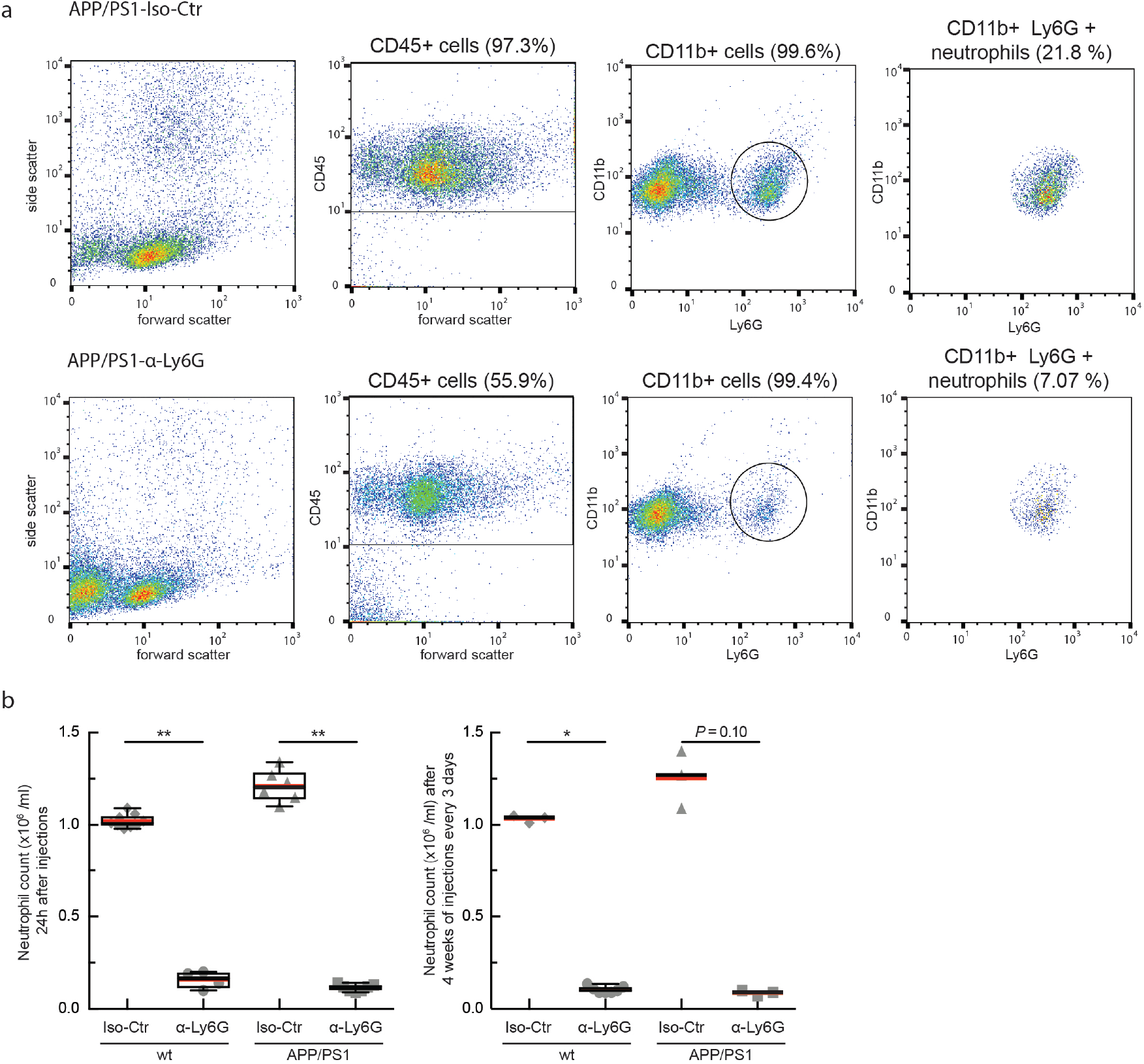
Treatment with α-Ly6G leads to neutrophil depletion in both APP/PS1 and wildtype control mice. (a) Representative flow cytometry data for blood drawn from APP/PS1 mice 24 hours after treatment with isotype control antibodies (top row) and α-Ly6G (bottom row). Left column shows forward and side scattering from entire population of blood cells (after lysing and removing red blood cells). The second column shows the gate on CD45+ cells, indicating leukocytes. The third column shows expression of CD11b (high for monocytes and neutrophils) and Ly6G (high for neutrophils) for the CD45+ cells. Cells with high expression levels of both CD11b and Ly6G were considered to be neutrophils (right column). (b) Neutrophil counts for APP/PS1 and wt mice 24 hr after treatment with α-Ly6G or isotype control antibodies (left) and after 4 weeks of treatment every three days (right) (24 hr data: wt Iso-Ctr: n=9 mice; wt Ly6G: n=4; APP/PS1 Iso-Ctr: n=6; APP/PS1 Ly6G: n=7; 4 week data: wt Iso-Ctr: n=3; wt Ly6G: n=7; APP/PS1 Iso-Ctr: n=3; APP/PS1 Ly6G: n=3; *p<0.05, **p<0.01, Mann-Whitney comparison between Iso-Ctr and Ly6G treated animals).

**Extended Data Fig. 15.**
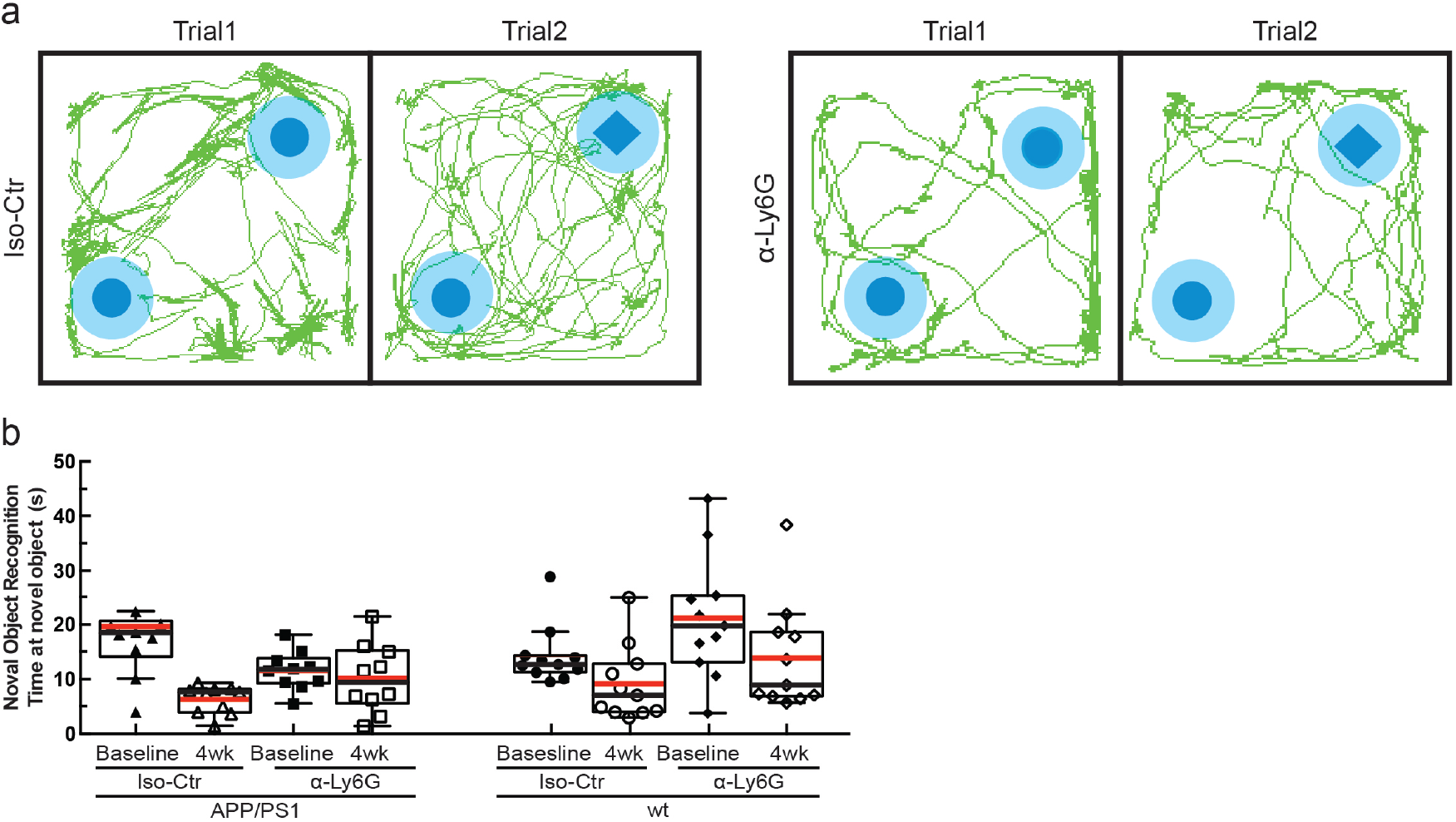
Representative map of animal location and time spent at the novel object in wild type controls and APP/PS1 animals treated with α-Ly6G or isotype control antibodies. (a) Tracking of mouse nose location from video recording during training and trial phases of novel object recognition task taken 4 weeks after administration of α-Ly6G or isotype control antibodies every three days in APP/PS1 mice. (b) Time spent at the novel object (APP/PS1 Iso-Ctr: 10 mice; APP/PS1 α-Ly6G: 10 mice; wt Iso-Ctr: 11 mice; wt α-Ly6G: 11 mice; no significant differences among groups as determined by Kruskal-Wallis one-way ANOVA).

**Extended Data Fig. 16.**
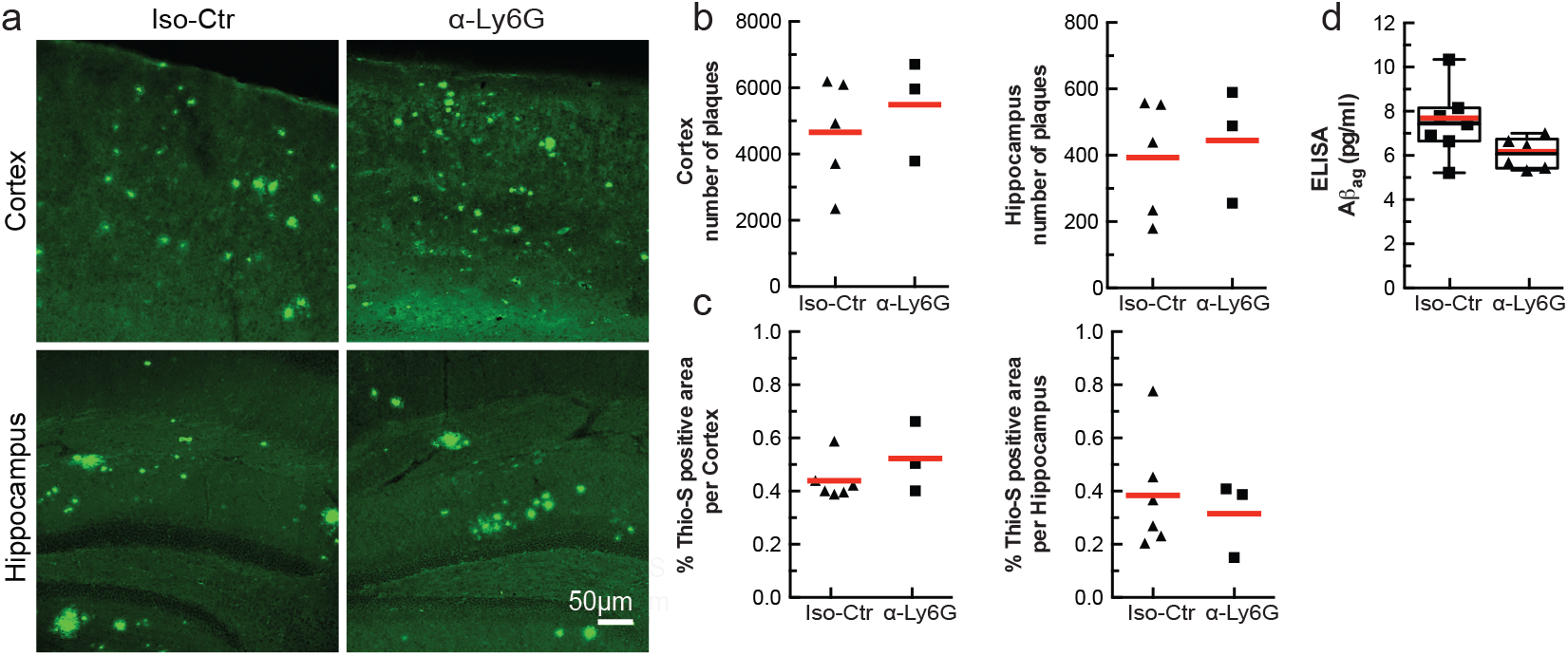
Amyloid plaque density and concentration of amyloid-beta oligomers were not changed in 11-month-old APP/PS1 animals treated with α-Ly6G every three days for a month. (a) Thioflavin-S staining of amyloid plaques in representative cortical sections (upper 2 panels) and hippocampal sections (lower 2 panels) for APP/PS1 mice treated with isotype control antibodies (left panels) or α-Ly6G (right panels). (b) Number of amyloid plaques in the cortex (left) and hippocampus (right) for APP/PS1 mice after one month of treatment (Iso-Ctr: 5 mice; α-Ly6G: 3 mice). (c) Percentage of tissue section positive for Thioflavin-S in the cortex (left) and hippocampus (right) (Iso-Ctr: 6 mice; α-Ly6G: 3 mice). (d) ELISA measurements of Aβ aggregate concentrations after 4 weeks of treatment with α-Ly6G or isotype control antibodies every three days (Iso-Ctr: 7 mice; α-Ly6G: 6 mice).

**Extended Data Fig. 17.**
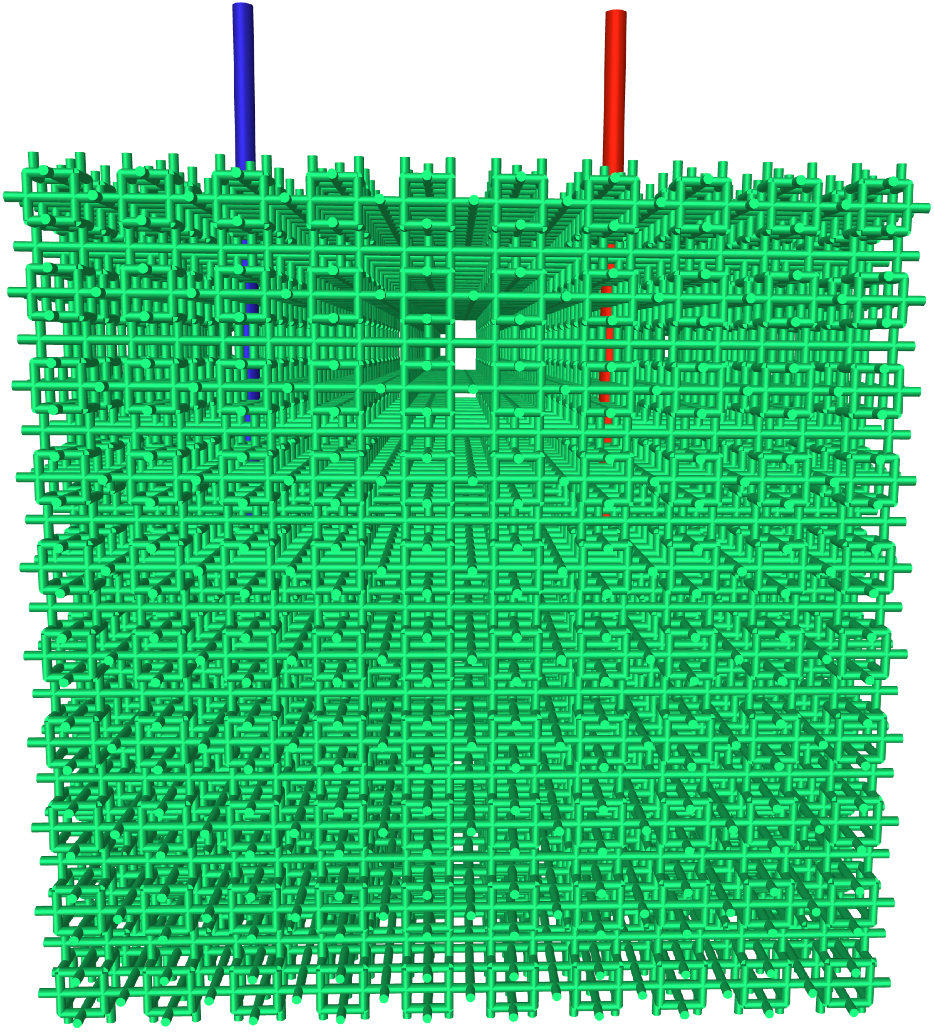
Synthetic capillary network of order three. Capillaries are indicated in green, while red and blue indicate the single feeding arteriole and draining venule, respectively.

**Extended Data Fig. 18.**
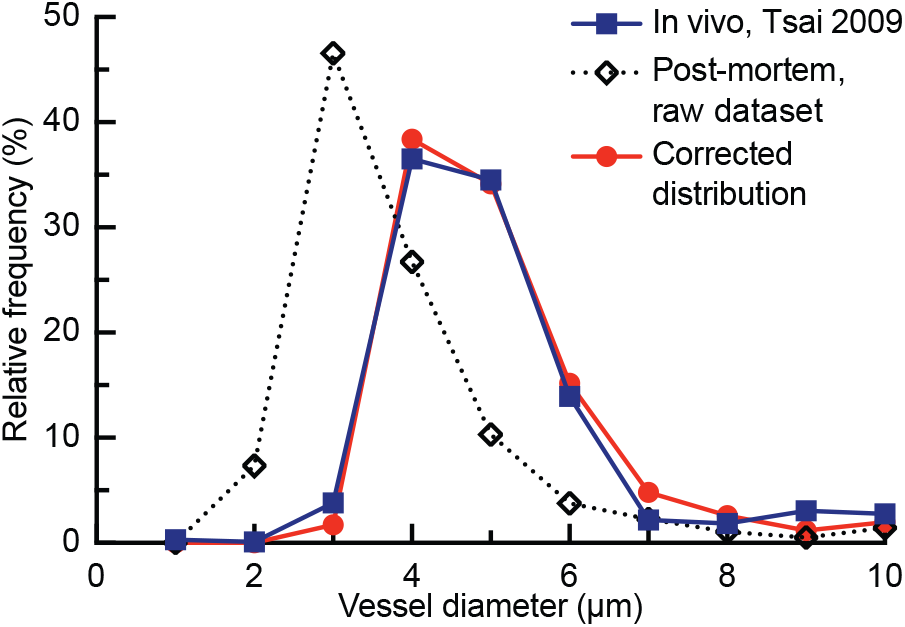
Histogram of mouse capillary diameters from *in vivo* measurements and post-mortem vascular casts. The diameter correction described in Eq. 3 closely aligned the post mortem diameters to the *in vivo* data.

**Extended Data Fig. 19.**
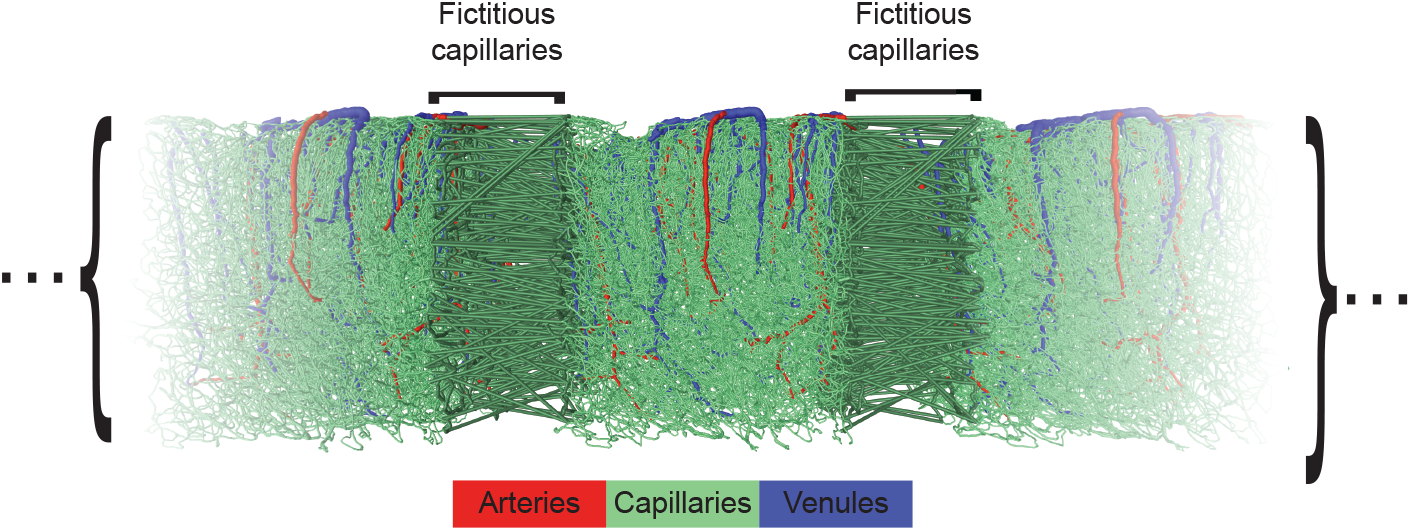
Illustration of the pseudo-periodic boundary conditions. Vessels categorized as arterioles are labeled in red, venules in blue, and capillaries in green.

**Extended Data Fig. 20.**
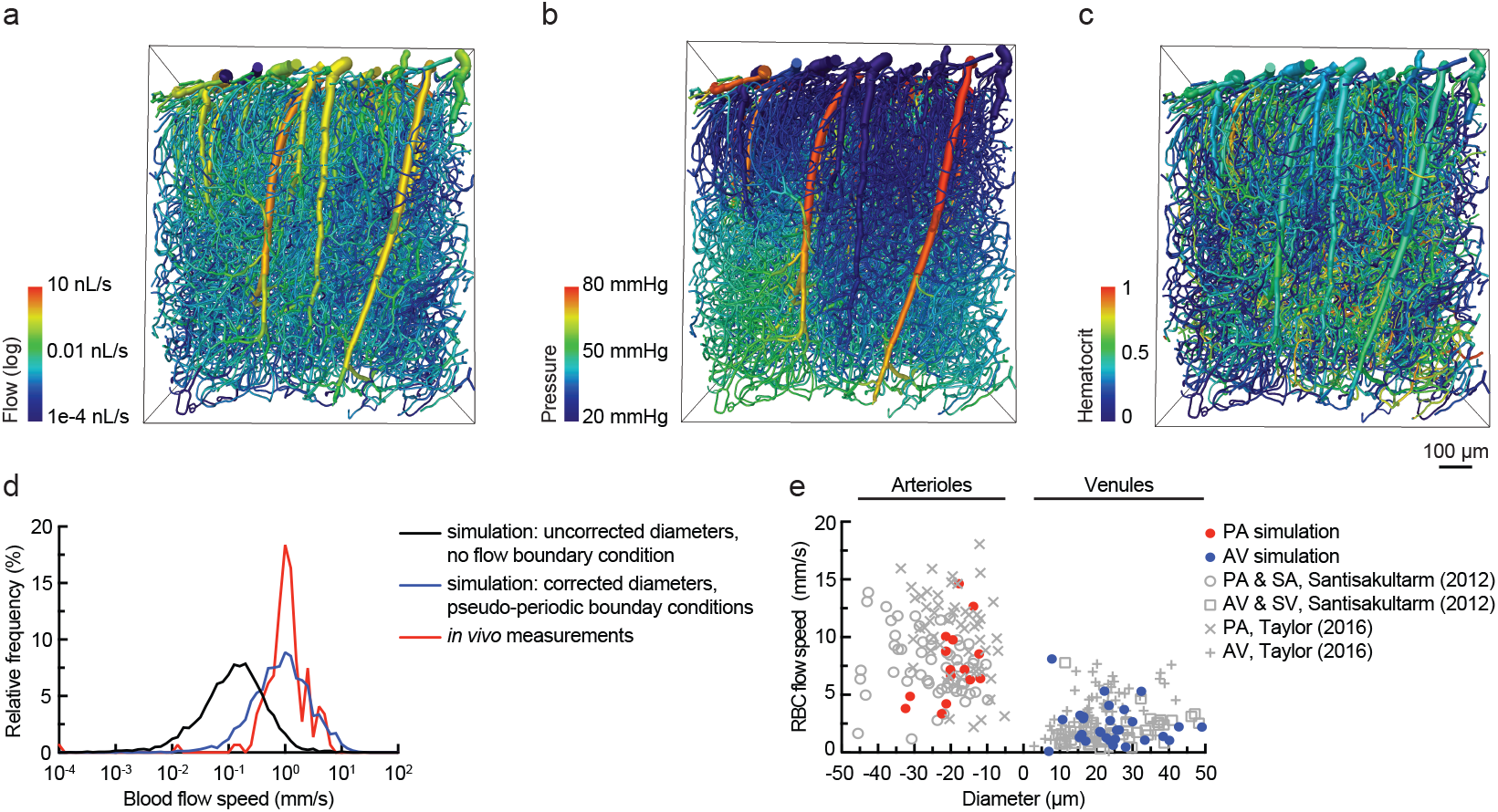
Validation of simulations. Spatial distribution of simulated blood flow (a), pressure (b), and hematocrit (c) in each vessel in the mouse vascular network. (d) Comparison of red blood cell velocities in capillaries in the top 300-μm of mouse cortex from experimental, *in vivo* measurements (red line), simulations with pseudo-periodic boundary conditions with corrected diameters (blue line), and no-flow boundary conditions without corrected diameters (black line). (e) Relationship between red blood cell speed and vessel diameter in arterioles and venules in calculations (solid red and blue dots) and experimental measurements (grey points).

**Extended Data Fig. 21.**
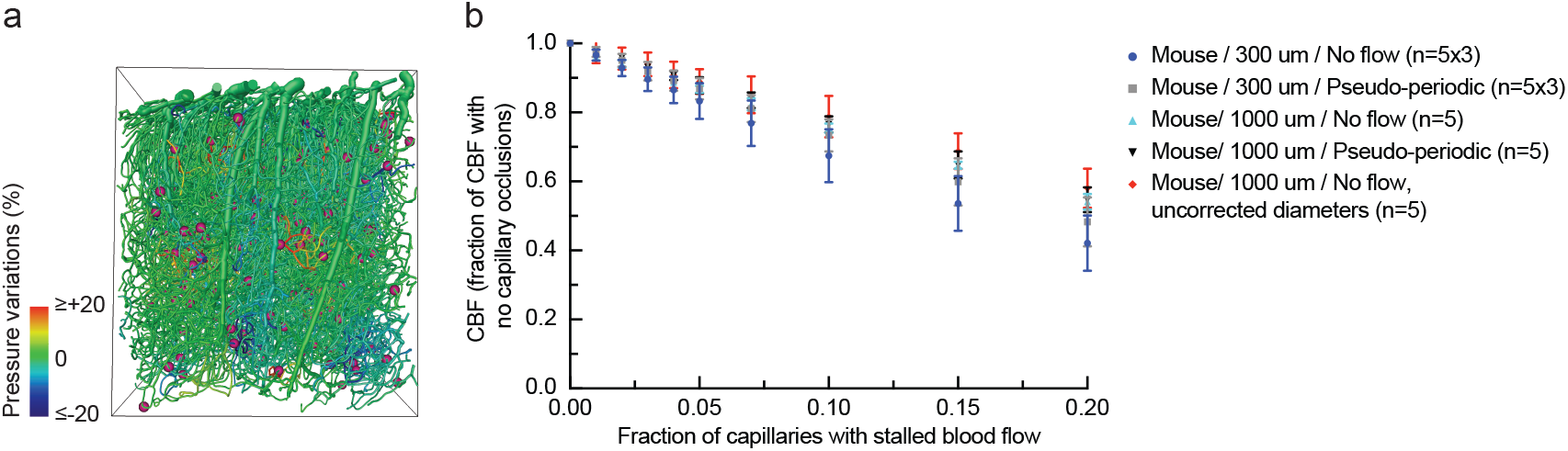
Calculated blood flow decreases due to capillary stalls was robust with respect to simulation parameters. (a) Pressure changes in mouse cortical vessel network due to randomly placed occlusions in 2% of capillaries. The corresponding flow changes are shown in Fig. 1J. (b) Calculated flow changes due to the occlusion of varying proportions of the capillaries using the full mouse dataset (100ü μm) or truncated datasets (1000x300 μm) with periodic or no flow boundary conditions, and with or without corrected vessel diameters. Error bars represent SD across n independent simulations (whole domain: n=5; 300 μm slices: n=5 for each of 3 slices).

### Supplementary Table

**Table S1:** Group sizes, statistical tests, and notes for main Figure panels.

| **Figure panel** | **Groups compared** | **Statistical tests and Notes** |
| --- | --- | --- |
| **1C, D** | APP/PS1: 28 mice,<br>~22,400 capillaries<br>wt: 12 mice,<br>~9,600 capillaries | Mann-Whitney, ****p<0.0001<br>Each data point represents one mouse in which > 800 capillaries were scored as flowing or stalled.<br>The lines in panel D represent a sliding average with a 10-week window and the shaded areas represent 95% confidence intervals. |
| **1F** | APP/PS1: 7 mice<br>Stalled: n ~ 120<br>Flowing: n = ~ 8,700 |  |
| **1G** | APP/PS1: 7 mice<br>Stalled: n ~ 120<br>Flowing: n = ~ 8,600 |  |
| **1J** |  | Error bars represent SD across for five independent simulations. |
| **2B** | APP/PS1: 10 mice,<br>235 stalled capillaries | Error bars represent 95% confidence intervals. |
| **2C** | Stalled: n = 116<br>Flowing: n = 8,431 | Mann-Whitney, ****p<0.0001 |
| **2D** | APP/PS1: 7 mice<br>Stalled: n ~ 120 |  |
|  | Flowing: n ~ 9,000 |  |
| **2E** | APP/PS1: 3 mice,<br>31 capillaries |  |
| **2G** | APP/PS1: 4 mice,<br>49 stalled capillaries<br>followed from first<br>imaging session |  |
| **3A** | α-Ly6G: 6 mice,<br>~4,800 capillaries<br>Iso-Ctr: 6 mice,<br>~4,800 capillaries | Mann-Whitney, **p<0.01 |
| **3C** | young APP/PS1 Iso-Ctr: 5 mice, 32<br>arterioles<br>old APP/PS1 Iso-Ctr: 3 mice, 18<br>arterioles<br>young wt α-Ly6G: 5<br>mice, 30 arterioles<br>young APP/PS1 α-<br>Ly6G: 5 mice, 33<br>arterioles<br>old APP/PS1 α-<br>Ly6G: 3 mice, 22<br>arterioles | Kruskal-Wallis one-way ANOVA with post-hoc<br>using Dunn's multiple comparison correction,<br>**p<0.01 and ***p<0.001 |
| **3E** | wt α-Ly6G: 10 mice<br>APP/PS1 α-Ly6G: 10 | Ordinary one-way ANOVA with post hoc using<br>Tukey's multiple comparison correction to |
|  | mice<br>APP/PS1 Iso-Ctr: 10 mice | compare across groups, *p<0.05<br>Paired t-test to compare baseline and after treatment within a group, ++p<0.01<br>Each data point indicates a single mouse and lines connecting baseline and after measurements indicate the same animal. |
| **4C, D, E** | APP/PS1 Iso-Ctr: 10 mice<br>APP/PS1 α-Ly6G: 10 mice<br>wt α-Ly6G: 11 mice<br>wt Iso-Ctr: 11 mice | Kruskal-Wallis one-way ANOVA with post-hoc using Dunn's multiple comparison correction to compare across groups, *p<0.05 and **p<0.01<br>Wilcoxon matched-pairs signed rank test relative to the baseline measurement to compare baseline and after treatment within a group, +p<0.05 and ++p<0.01 |
| **4F** | Iso-Ctr: 6 mice<br>α-Ly6G: 7 mice | Mann-Whitney<br>**p <0.01 |

## Supplementary Movies

**Movie S1. Two-photon image stacks of fluorescently labeled blood vessels from APP/PS1 mice.** Capillaries with stalled blood flow are indicated with red circles.

**Movie S2. Two-photon image stacks of fluorescently labeled blood vessels from wt mice.** Capillaries with stalled blood flow are indicated with red circles.

**Movie S3. Two-photon image stacks of fluorescently-labeled blood vessels APP/PS1 mouse when anesthetized.** Capillaries with stalled blood flow are indicated with red circles. Animal was anesthetized by breathing 1.5% isoflurane.

**Movie S4. Two-photon image stacks of fluorescently-labeled blood vessels from the same APP/PS1 mouse (Movie. S3) when awake.** Capillaries with stalled blood flow are indicated with red circles.

